# From recency to central tendency biases in working memory: a unifying network model

**DOI:** 10.1101/2022.05.16.491352

**Authors:** Vezha Boboeva, Alberto Pezzotta, Claudia Clopath, Athena Akrami

## Abstract

The central tendency bias, or contraction bias, is a phenomenon where the judgment of the magnitude of items held in working memory appears to be biased towards the average of past observations. It is assumed to be an optimal strategy by the brain, and commonly thought of as an expression of the brain’s ability to learn the statistical structure of sensory input. On the other hand, recency biases such as serial dependence are also commonly observed, and are thought to reflect the content of working memory. Recent results from an auditory delayed comparison task in rats, suggest that both biases may be more related than previously thought: when the posterior parietal cortex (PPC) was silenced, both short-term and contraction biases were reduced. By proposing a model of the circuit that may be involved in generating the behavior, we show that a volatile working memory content susceptible to shifting to the past sensory experience – producing short-term sensory history biases – naturally leads to contraction bias. The errors, occurring at the level of individual trials, are sampled from the full distribution of the stimuli, and are not due to a gradual shift of the memory towards the sensory distribution’s mean. Our results are consistent with a broad set of behavioral findings and provide predictions of performance across different stimulus distributions and timings, delay intervals, as well as neuronal dynamics in putative working memory areas. Finally, we validate our model by performing a set of human psychophysics experiments of an auditory parametric working memory task.

## Introduction

A fundamental question in neuroscience relates to how brains efficiently process the statistical regularities of the environment to guide behavior. Exploiting such regularities may be of great value to survival in the natural environment, but may lead to biases in laboratory tasks. Repeatedly observed across species and sensory modalities is the central tendency (“contraction”) bias, where performance in perceptual tasks seemingly reflects a shift of the working memory representation towards the mean of the sensory history [1–6]. Equally common are sequential biases, either attractive or repulsive, towards the immediate sensory history [7, 5, 8–14, 6, 15].

It is commonly thought that these biases occur due to disparate mechanisms - contraction bias is widely thought to be a result of learning the statistical structure of the environment, whereas serial biases are thought to reflect the contents of working memory [16, 17]. Recent evidence, however, challenges this picture: our recent study of a parametric working memory task discovered that the rat posterior parietal cortex (PPC) plays a key role in modulating contraction bias [7]. When the region is optogenetically inactivated, contraction bias is attenuated. Intriguingly, however, this is also accompanied by the suppression of bias effects induced by the recent history of the stimuli, suggesting that the two phenomena may be interrelated. Interestingly, other behavioral components, including working memory of immediate sensory stimuli (in the current trial), remain intact. In another recent study with humans, a double dissociation was reported between three cohorts of subjects: subjects on the autistic spectrum (ASD) expressed reduced biases due to recent statistics, whereas dyslexic subjects (DYS) expressed reduced biases towards long-term statistics, relative to neurotypical subjects (NT) [16]. Finally, further complicating the picture is the observation of not only attractive serial dependency, but also repulsive biases [18]. It is as of yet unclear how such biases occur and what mechanisms underlie such history dependencies.

These findings stimulate the question of whether contraction bias and the different types of serial biases are actually related, and if so, how. Although normative models have been proposed to explain these effects [19, 18, 16], the neural mechanisms and circuits underlying them remain poorly understood. We address this question through a model of the putative circuit involved in giving rise to the behavior observed in [7]. Our model consists of two continuous (bump) attractor sub-networks, representing a working memory (WM) module and the PPC. Given the finding that PPC neurons carry more information about stimuli presented during previous trials, the PPC module integrates inputs over a longer timescale relative to the WM network, and incorporates firing rate adaptation.

We find that both contraction bias and short-term sensory history effects emerge in the WM network as a result of inputs from the PPC network. Importantly, we see that these effects do not necessarily occur due to separate mechanisms. Rather, in our model, contraction bias emerges as a statistical effect of errors in working memory, occurring due to the persisting memory of stimuli shown in the preceding trials. The integration of this persisting memory in the WM module competes with that of the stimulus in the current trial, giving rise to short-term history effects. We conclude that contraction biases in such paradigms may not necessarily reflect explicit learning of regularities or an “attraction towards the mean”, on individual trials. Rather, it may be an effect emerging at the level of average performance, when in each trial, errors are made according to the recent sensory experiences whose distribution follow that of the input stimuli. Furthermore, using the same model, we also show that the biases towards long-term (short-term) statistics inferred from the performance of ASD (DYS) subjects [16] may actually reflect short-term biases extending more or less into the past with respect to NT subjects, challenging the hypothesis of a double-dissociation mechanism. Last, we show that as a result of neuronal integration of inputs and adaptation, in addition to attraction effects occurring on a short timescale, repulsion effects are observed on a longer timescale [18].

We make specific predictions on neuronal dynamics in the PPC and downstream working memory areas, as well as how contraction bias may be altered, upon manipulations of the sensory stimulus distribution, inter-trial and inter-stimulus delay intervals. We show agreements between the model and our previous results in humans and rats. Finally, we test our model predictions by performing new human auditory parametric working memory tasks. The data is in agreement with our model and not with an aternative Bayesian model.

## 1 Results

### 1.1 The PPC as a slower integrator network

Parametric working memory (PWM) tasks involve the sequential comparison of two graded stimuli that differ along a physical dimension and are separated by a delay interval of a few seconds (Fig. 1 A and B) [20, 7, 19]. A key feature emerging from these studies is contraction bias, where the averaged performance is as if the memory of the first stimulus progressively shifts towards the center of a prior distribution built from past sensory history (Fig. 1 C). Additionally, biases towards the most recent sensory stimuli (immediately preceding trials) have also been documented [7, 5].

**Figure 1:**
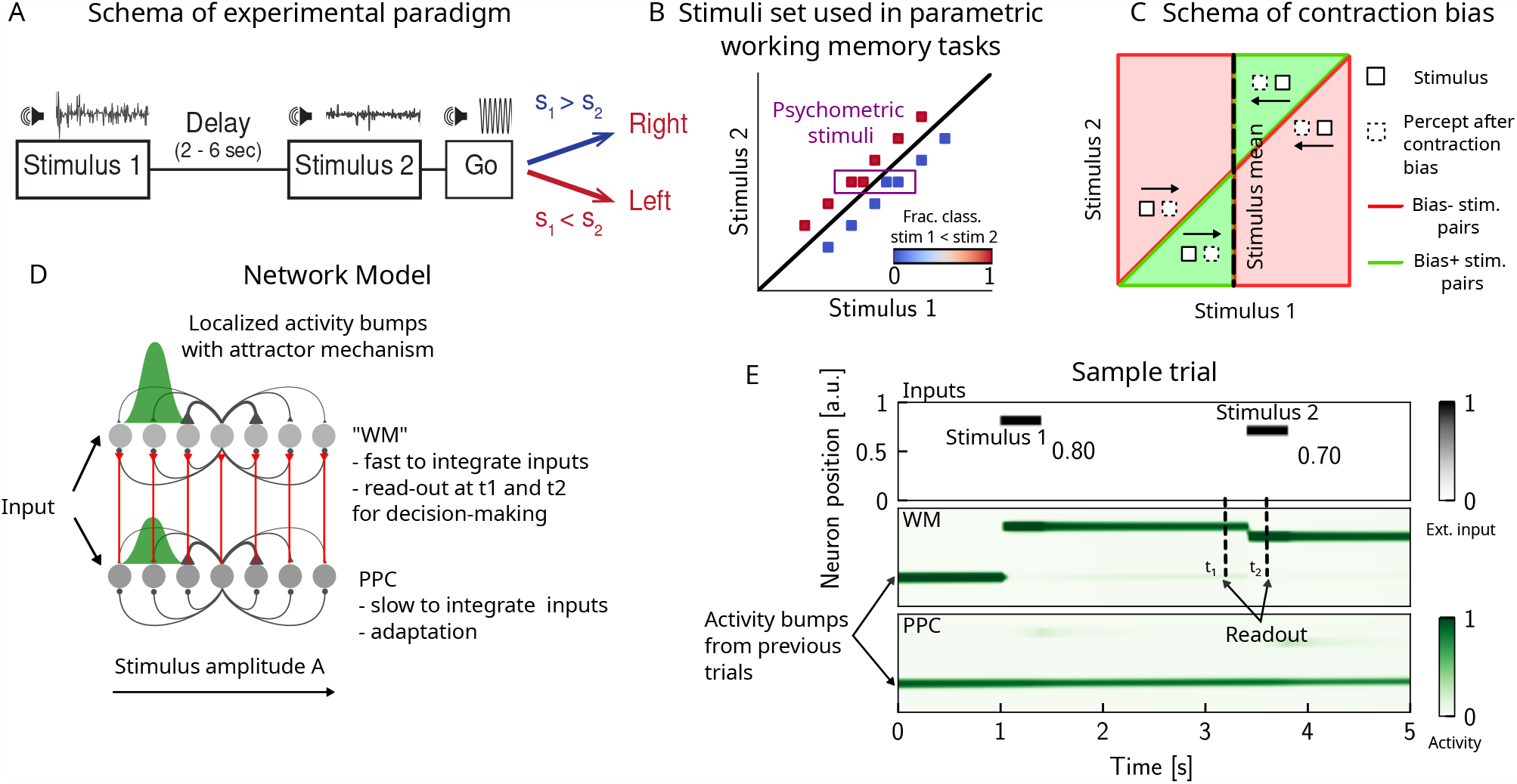
The PPC as a slower integrator network. **(A)** In any given trial, a pair of stimuli (here, sounds) separated by a variable delay interval is presented to a subject. After the second stimulus, and after a go cue, the subject must decide which of the two sounds is louder by pressing a key (humans) or nose-poking in an appropriate port (rats). **(B)** The stimulus set. The stimuli are linearly separable, and stimulus pairs are equally distant from the *s*_1_ = *s*_2_ diagonal. Error-free performance corresponds to network dynamics from which it is possible to classify all the stimuli below the diagonal as *s*_1_ *> s*_2_ (shown in blue) and all stimuli above the diagonal as *s*_1_ *< s*_2_ (shown in red). An example of a correct trial can be seen in (E). In order to assay the psychometric threshold, several additional pairs of stimuli are included (purple box), where the distance to the diagonal *s*_1_ = *s*_2_ is systematically changed. The colorbar expresses the fraction classified as *s*_1_ *< s*_2_. **(C)** Schematics of contraction bias in delayed comparison tasks. Performance is a function of the difference between the two stimuli, and is impacted by contraction bias, where the base stimulus *s*_1_ is perceived as closer to the mean stimulus. This leads to a better/worse (green/red area) performance, depending on whether this “attraction” increases (Bias+) or decreases (Bias-) the discrimination between the base stimulus *s*_1_ and the comparison stimulus *s*_2_. **(D)** Our model is composed of two modules, representing working memory (WM), and sensory history (PPC). Each module is a continuous one-dimensional attractor network. Both networks are identical except for the timescales over which they integrate external inputs; PPC has a significantly longer integration timescale and its neurons are additionally equipped with neuronal adaptation. The neurons in the WM network receive input from those in the PPC, through connections (red lines) between neurons coding for the same stimulus. Neurons (gray dots) are arranged according to their preferential firing locations. The excitatory recurrent connections between neurons in each network are a symmetric, decreasing function of their preferential firing locations, whereas the inhibitory connections are uniform (black lines). For simplicity, connections are shown for a single pre-synaptic neuron (where there is a bump in green). When a sufficient amount of input is given to a network, a bump of activity is formed, and sustained in the network when the external input is subsequently removed. This activity in the WM network is read out at two time points: slightly before and after the onset of the second stimulus, and is used to assess performance. **(E)** The task involves the comparison of two sequentially presented stimuli, separated by a delay interval (top panel, black lines). The WM network integrates and responds to inputs quickly (middle panel), while the PPC network integrates inputs more slowly (bottom panel). As a result, external inputs (corresponding to stimulus 1 and 2) are enough to displace the bump of activity in the WM network, but not in the PPC. Instead, inputs coming from the PPC into the WM network are not sufficient to displace the activity bump, and the trial is consequently classified as correct. In the PPC, instead, the activity bump corresponds to a stimulus shown in previous trials.

In order to investigate the circuit mechanisms by which such biases may occur, we use two identical one-dimensional continuous attractor networks to model WM and PPC modules. Neurons are arranged according to their preferential firing locations in a continuous stimulus space, representing the amplitude of auditory stimuli. Excitatory recurrent connections between neurons are symmetric and a monotonically decreasing function of the distance between the preferential firing fields of neurons, allowing neurons to mutually excite one another; inhibition, instead, is uniform. Together, both allow a localized bump of activity to form and be sustained (Fig. 1 D and E) [21–29]. Both networks have free boundary conditions. Neurons in the WM network receive inputs from neurons in the PPC coding for the same stimulus amplitude (Fig. 1 D). Building on experimental findings [30–35], we designed the PPC network such that it integrates activity over a longer timescale compared to the WM network (Sect. 3.1). Moreover, neurons in the PPC are equipped with neural adaptation, that can be thought of as a threshold that dynamically follows the activation of a neuron over a longer timescale.

To simulate the parametric WM task, at the beginning of each trial, the network is provided with a stimulus *s*_1_ for a short time via an external current *I*_ext_ as input to a set of neurons (see Tab. 1). Following *s*_1_, after a delay interval, a second stimulus *s*_2_ is presented (Fig. 1 E). The pair (*s*_1_, *s*_2_) is drawn from the stimulus set shown in Fig. 1 B, where they are all equally distant from the diagonal *s*_1_ = *s*_2_, and are therefore of equal nominal discrimination, or difficulty. The stimuli (*s*_1_, *s*_2_) are co-varied in each trial so that the task cannot be solved by relying on only one of the stimuli [36]. As in the study in Ref. [7] using an interleaved design, across consecutive trials, the inter-stimulus delay intervals are randomized and sampled uniformly between 2, 6 and 10 seconds. The inter-trial interval, instead, is fixed at 5 seconds.

**Table 1:**
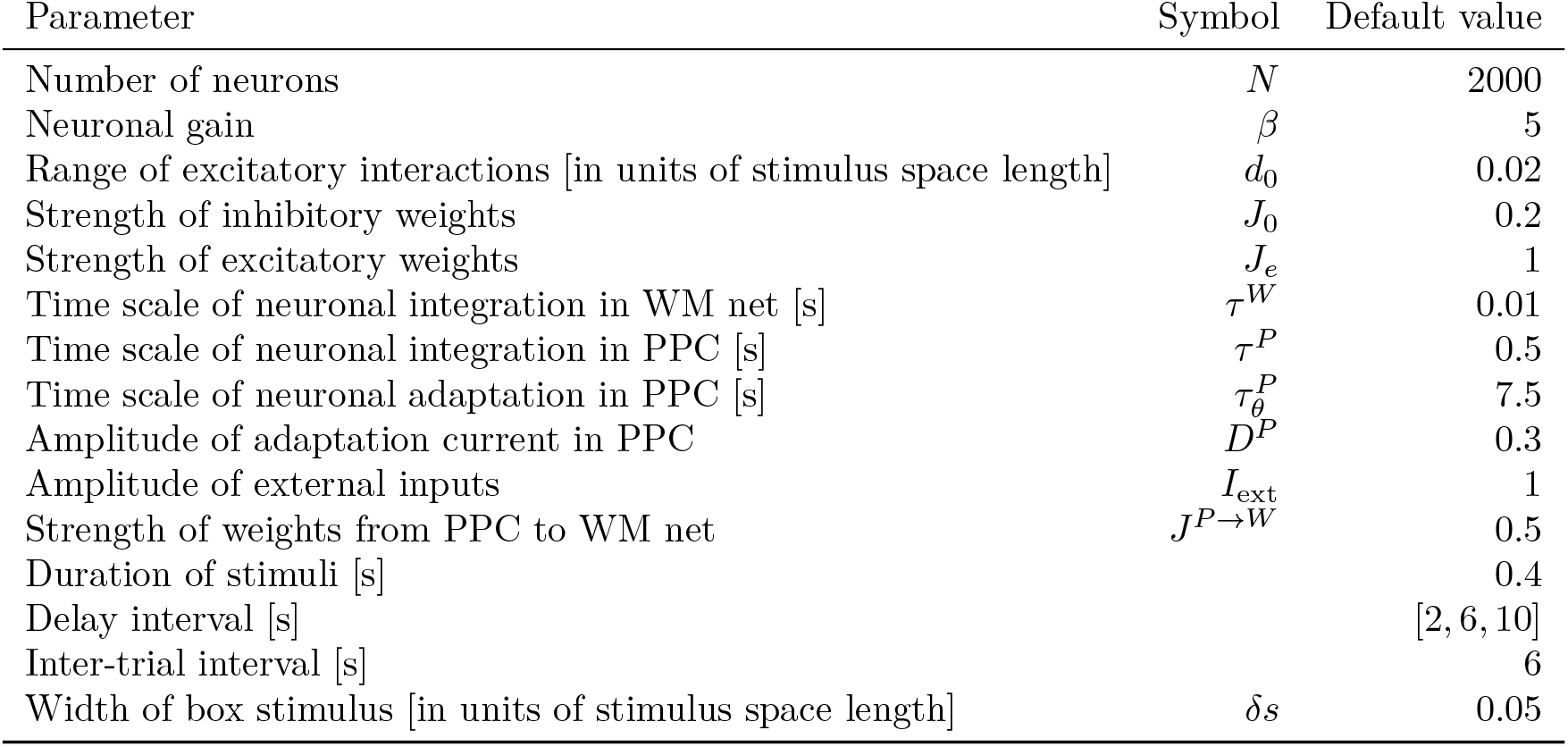
Simulation parameters, when not explicitly mentioned. Used to produce Figs. 1, 2, 3, 4, 5, 8, S2, S3, S5.

We additionally include psychometric pairs (indicated in the box in Fig. 1 B) where the distance to the diagonal, hence the discrimination difficulty, is varied. The task is a binary comparison task that aims at classifying whether *s*_1_ *< s*_2_ or vice-versa. In order to solve the task, we record the activity of the WM network at two time points: slightly before and after the onset of *s*_2_ (Fig. 1 E). We repeat this procedure across many different trials, and use the recorded activity to assess performance (see (Sect. 3.2) for details). Importantly, at the end of each trial, the activity of both networks is not re-initialized, and the state of the network at the end of the trial serves as the initial network configuration for the next trial.

### 1.2 Contraction bias and short-term stimulus history effects as a result of PPC network activity

Probing the WM network performance on psychometric stimuli (Fig. 1 B, purple box, 10% of all trials) shows that the comparison behavior is not error-free, and that the psychometric curves (different colors) differ from the optimal step function (Fig. 2 A, green dashed line). The performance on pychometric trials is also better for shorter inter-stimulus delay intervals, as has been shown in previous work [37, 7]. In our model, errors are caused by the displacement of the activity bump in the WM network, due to the inputs from the PPC network. These displacements in the WM activity bump can result in different outcomes: by displacing it *away* from the second stimulus, they either do not affect the performance or improve it (Fig. 2 B right panel, “Bias+”), if noise is present. Conversely, the performance can suffer, if the displacement of the activity bump is *towards* the second stimulus (Fig. 2 B left panel, “Bias-”). Note, however, that in these two specific trials, the activity bump in PPC is strong, and it displaces the activity bump in the WM network, but this is not the only kind of dynamics present in the network (see Sect.1.3 for a more detailed analysis of the network dynamics).

**Figure 2:**
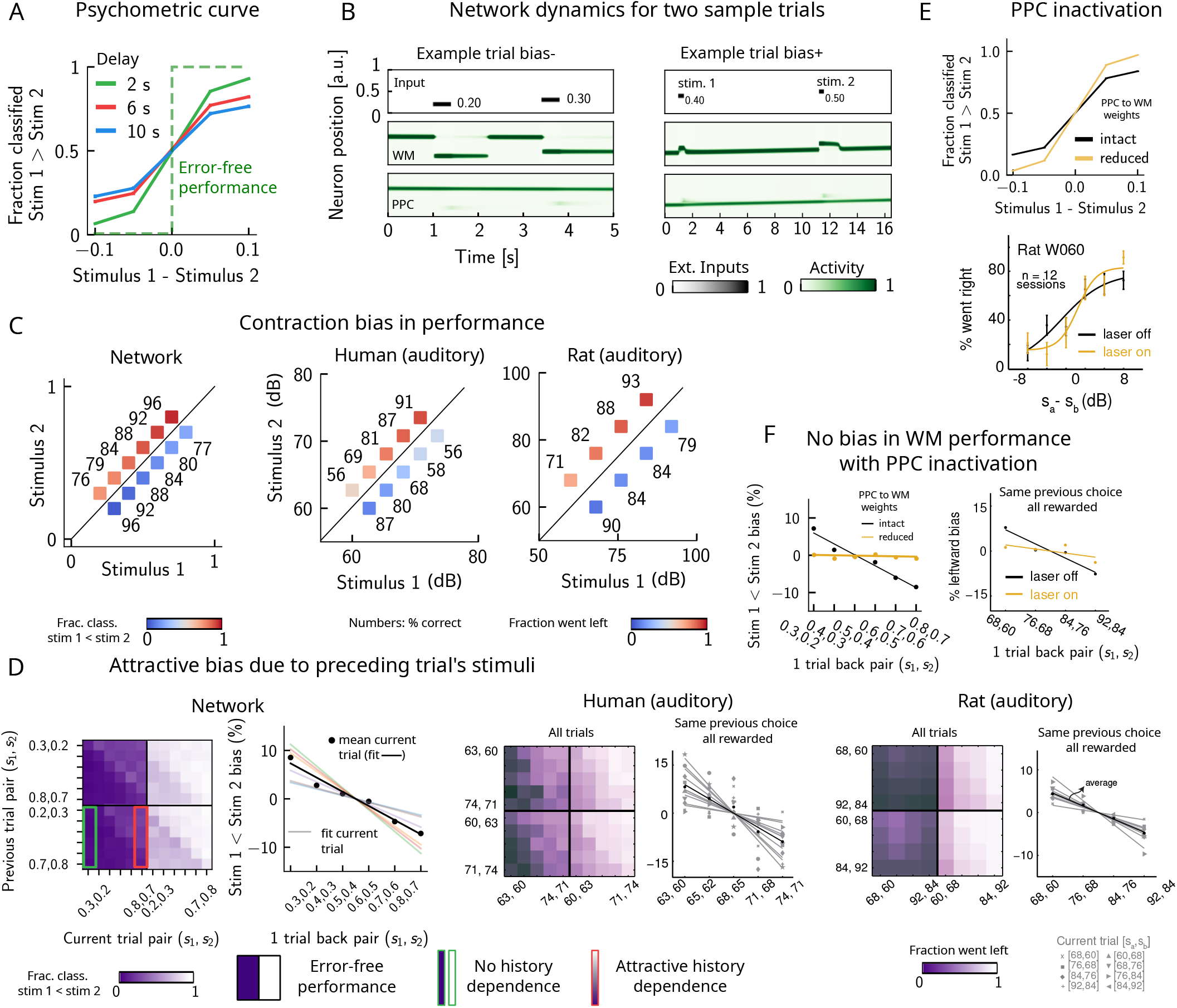
Contraction bias and short-term sensory history effects as a result of PPC network activity. **(A)** Performance of network model for psychometric stimuli (colored lines) is not error-free (green dashed lines). A shorter inter-stimulus delay interval yields a better performance. **(B)** Errors occur due to the displacement of the bump representing the first stimulus *s*_1_ in the WM network. Depending on the direction of this displacement with respect to *s*_2_, this can give rise to trials in which the comparison task becomes harder (easier), leading to negative (positive) biases (top and bottom panels). Top sub-panel: stimuli presented to both networks in time. Middle/ bottom sub-panels show activity of WM and PPC networks (in green). **(C)** Left: performance is affected by contraction bias – a gradual accumulation of errors for stimuli below (above) the diagonal upon increasing (decreasing) *s*_1_. Colorbar indicates fraction of trials classified as *s*_1_ *< s*_2_. Middle and Right: for comparison, data from the auditory version of the task performed in humans and rats. Data from Ref. [7]. **(D)** Panel 1: For each combination of current (x-axis) and previous trial’s stimulus pair (y-axis), fraction of trials classified as *s*_1_ *< s*_2_ (colorbar). Performance is affected by preceding trial’s stimulus pair (modulation along the y-axis). For readability, only some tick-labels are shown. Panel 2: bias, quantifying the (attractive) effect of previous stimulus pairs. Colored lines correspond to linear fits of this bias for each pair of stimuli in the current trial. Black dots correspond to average over all current stimuli, and black line is a linear fit. These history effects are attractive: the smaller the previous stimulus, the higher the probability of classifying the first stimulus of the current trial *s*_1_ as small, and vice-versa. Panel 3: human auditory trials. Percentage of trials in which humans chose left for each combination of current and previous stimuli; vertical modulation indicates attractive effect of preceding trial. Panel 4: Percentage of trials in which humans chose left minus the average value of left choices, as a function of the stimuli of the previous trial, for fixed previous trial response choice and reward. Panel 5 and 6: same as panels 3 and 4 but with rat auditory trials. Data from Ref. [7]. **(E)** Top: performance of network, when the weights from the PPC to the WM network is weakened, is improved for psychometric stimuli (yellow curve), relative to the intact network (black curve). Bottom: psychometric curves for rats (only shown for one rat) are closer to error-free during PPC inactivation (yellow) than during control trials (black). **(F)** Left: the attractive bias due to the effect of the previous trial is present with the default weights (black line), but is eliminated with reduced weights (yellow line). Right: while there is bias induced by previous stimuli in the control experiment (black), this bias is reduced under PPC inactivation (yellow). Experimental figures reproduced with permission from Ref.[7].

Performance on stimulus pairs that are equally distant from the *s*_1_ = *s*_2_ diagonal can be similarly impacted and the network produces a pattern of errors that is consistent with contraction bias: performance is at its minimum for stimulus pairs in which *s*_1_ is either largest or smallest, and at its maximum for stimulus pairs in which *s*_2_ is largest or smallest (Fig. 2 C, left panel) [19, 38, 7, 39, 40]. These results are consistent with the performance of humans and rats on the auditory task, as previously reported (Fig. 2 C, middle and right panels, data from Akrami et al 2018 [7]).

Can the same circuit also give rise to short-term sensory history biases [7, 41]? We analyzed the fraction of trials the network response was “*s*_1_ *< s*_2_” in the current trial conditioned on stimulus pairs presented in the previous trial, and we found that the network behavior is indeed modulated by the preceding trial’s stimulus pairs (Fig. 2 D, panel 1). We quantified these history effects as well as how many trials back they extend to. We computed the bias by plotting, for each particular pair (of stimuli) presented at the current trial, the fraction of trials the network response was “*s*_1_ *< s*_2_” as a function of the pair presented in the previous trial minus the mean performance over all previous trial pairs (Fig. 2 D, panel 2) [7]. Independent of the current trial, the previous trial exerts an “attractive” effect, expressed by the negative slope of the line: when the previous pair of stimuli is small, *s*_1_ in the current trial is, on average, misclassified as smaller than it actually is, giving rise to the attractive bias in the comparison performance; the converse holds true when the previous pair of stimuli happens to be large. These effects extend to two trials back, and are consistent with the performance of humans and rats on the auditory task (Fig. 2 D, panels 3-6, data from Akrami et al 2018 [7]).

It has been shown that inactivating the PPC, in rats performing the auditory delayed comparison task, markedly reduces the magnitude of contraction bias, without impacting non-sensory biases [7]. We assay the causal role of the PPC in generating the sensory history effects as well as contraction bias by weakening the connections from the PPC to the WM network, mimicking the inactivation of the PPC. In this case, we see that the performance for the psychometric stimuli is greatly improved (yellow curve, Fig. 2 E, top panel), consistent also with the inactivation of the PPC in rodents (yellow curve, Fig. 2 E, bottom panel, data from Akrami et al 2018 [7]). Performance is improved also for all pairs of stimuli in the stimulus set (Fig. S3 A). The breakdown of the network response in the current trial conditioned on the specific stimulus pair preceding it reveals that the previous trial no longer exerts a notable modulating effect on the current trial (Fig. S3 B). Quantifying this bias by subtracting the mean performance over all of the previous pairs reveals that the attractive bias is virtually eliminated (yellow curve, Fig. 2 F, left panel), consistent with findings in rats (Fig. 2 F, right panel, data from Akrami et al 2018 [7]).

Together, our results suggest a possible circuit through which both contraction bias and short-term history effects in a parametric working memory task may arise. The main features of our model are two continuous attractor networks, both integrating the same external inputs, but operating over different timescales. Crucially, the slower one, a model of the PPC, includes neuronal adaptation, and provides input to the faster one, intended as a WM circuit. In the next section, we show how the slow integration and firing rate adaptation in the PPC network give rise to the observed effects of sensory history.

### 1.3 Multiple timescales at the core of short-term sensory history effects

The activity bumps in the PPC and WM networks undergo different dynamics, due to the different timescales with which they integrate inputs, the presence of adaptation in the PPC, and the presence of global inhibition. The WM network integrates inputs over a shorter timescale, and therefore the activity bump follows the external input with high fidelity (Fig. 3 A (purple bumps) and B (purple line)). The PPC network, instead, has a longer integration timescale, and therefore fails to sufficiently integrate the input to induce a displacement of the bump to the location of a new stimulus, at each single trial. This is mainly due to the competition between the inputs from the recurrent connections sustaining the bump, and the external stimuli that are integrated elsewhere: if the former is stronger, the bump is not displaced. If, however, these inputs are weaker, they will not displace it, but may still exert a weakening effect via the global inhibition in the connectivity. The external input, as well as the presence of adaptation (Fig. S1 B and C) induce a small continuous drift of the activity bump that is already present from the previous trials (lower right panel of Fig. 2 B, Fig. 3 A (pink bumps) and B (pink line)). The build-up of adaptation in the PPC network, combined with the global inhibition from other neurons receiving external inputs, can extinguish the bump in that location (see also Fig. S1 for more details). Following this, the PPC network can make a transition to an incoming stimulus position (that may be either *s*_1_ or *s*_2_), and a new bump is formed. The resulting dynamics in the PPC are a mixture of slow drift over a few trials, followed by occasional jumps (Fig. 3 A).

**Figure 3:**
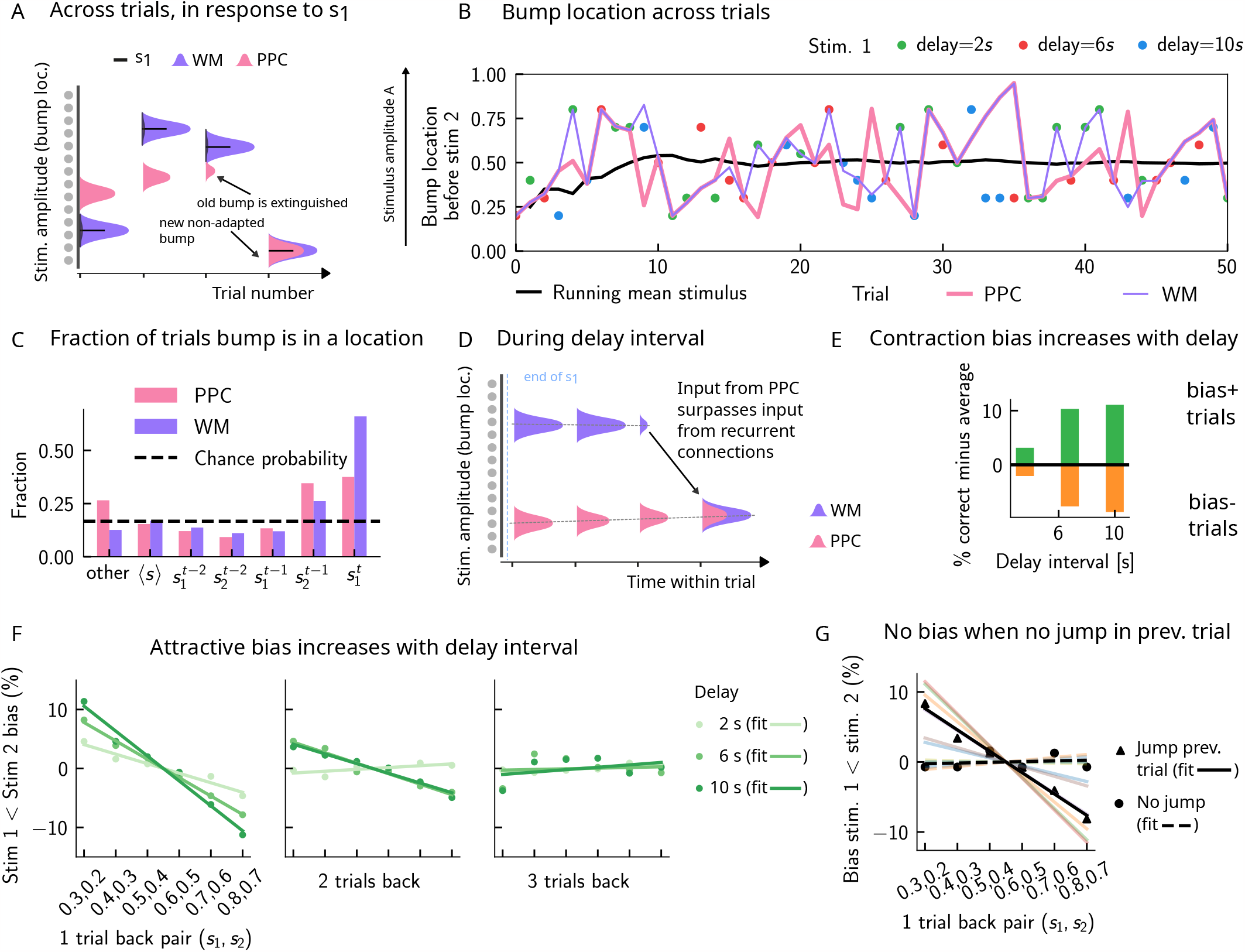
Multiple timescales at the core of short-term sensory history effects. **(A)** Schematics of activity bump dynamics in the WM vs PPC network. Whereas the WM responds quickly to external inputs, the bump in the PPC drifts slowly and adapts, until it is extinguished and a new bump forms. **(B)** The location of the activity bump in both the PPC (pink line) and the WM (purple line) networks, immediately before the onset of the second stimulus *s*_2_ of each trial. This location corresponds to the amplitude of the stimulus being encoded. The bump in the WM network closely represents the stimulus *s*_1_ (shown in colored dots, each color corresponding to a different delay interval). The PPC network, instead, being slower to integrate inputs, displays a continuous drift of the activity bump across a few trials, before it jumps to a new stimulus location, due to the combined effect of inhibition from incoming inputs and adaptation that extinguishes previous activity. **(C)** Fraction of trials in which the bump location corresponds to the base stimulus that has been presented 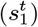 in the current trial, as well as the two preceding trials 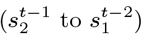. In the WM network, in the majority of trials, the bump coincides with the first stimulus of the current trial 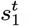. In a smaller fraction of the trials, it corresponds to the previous stimulus 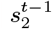, due to the input from the PPC. In the PPC network instead, a smaller fraction of trials consist of the activity bump coinciding with the current stimulus 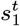. Relative to the WM network, the bump is more likely to coincide with the previous trial’s comparison stimulus 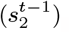. **(D)** During the inter-stimulus delay interval, in the absence of external sensory inputs, the activity bump in the WM network is mainly sustained endogenously by the recurrent inputs. It may, however, be destabilized by the continual integration of inputs from the PPC. **(E)** As a result, with an increasing delay interval, given that more errors are made, contraction bias increases. Green (orange) bars correspond to the performance in Bias+ (Bias-) regions, relative to the mean performance over all pairs (Fig. 1 C). **(F)** Left and middle: longer delay intervals allow for a longer integration times which in turn lead to a larger frequency of WM disruptions due to previous trials, leading to a larger previous-trial attractive biases (2*s* vs. 6*s* vs. 10*s*). Right: Weak repulsive effects for larger delays become apparent. Colored dots correspond to the bias computed for different values of the inter-stimulus delay interval, while colored lines correspond to their linear fits. **(G)** When neuronal adaptation is at its lowest in the PPC i.e. following a bump jump, the WM bump is maximally susceptible to inputs from the PPC. The attractive bias (towards previous stimuli) is present in trials in which the PPC network underwent a jump in the previous trial (black triangles, with black line a linear fit). Such biases are absent in trials where no jumps occur in the PPC in the previous trial (black dots, with dashed line a linear fit). Colored lines correspond to bias for specific pairs of stimuli in the current trial, regular lines for the jump condition, and dashed for the no jump condition.

As a result of such dynamics, relative to the WM network, the activity bump in the PPC represents the stimuli corresponding to the current trial in a smaller fraction of the trials, and represents stimuli presented in the previous trial in a larger fraction of the trials (Fig. 3 C). This yields short-term sensory history effects in our model (Fig. 2 D, and E), as input from the PPC lead to the displacement of the WM bump to other locations (Fig. 3 D). Given that neurons in the WM network integrate this input, once it has built up sufficiently, it can surpass the self-sustaining inputs from the recurrent connections in the WM network. The WM bump, then, can move to a new location, given by the position of the bump in the PPC (Fig. 3 D). As the input from the PPC builds up gradually, the probability of bump displacement in WM increases over time. This in return leads to an increased probability of contraction bias (Fig. 3 E), and short-term history (one-trial back) biases (Fig. 3 F), as the inter-stimulus delay interval increases.

Additionally, a non-adapted input from the PPC has a larger likelihood of displacing the WM bump. This is highest immediately following the formation of a new bump in the PPC, or in other words, following a “bump jump” (Fig. 3 F). As a result, one can reason that those trials immediately following a jump in the PPC are the ones that should yield the maximal bias towards stimuli presented in the previous trial. We therefore separated trials according to whether or not a jump has occurred in the PPC in the preceding trial (we define a jump to have occurred if the bump location across two consecutive trials in the PPC is displaced by an amount larger than the typical width of the bump (Sect. 3.1)). In line with this reasoning, only the set that included trials with jumps in the preceding trial yields a one-trial back bias (Fig. 3 G).

Removing neuronal adaptation entirely from the PPC network further corroborates this result. In this case, the network dynamics show a very different behavior: the activity bump in the PPC undergoes a smooth drift (Fig. S2 A), and the bump distribution is much more peaked around the mean (Fig. S2 B), relative to when adaptation is present (Fig. 4 A). In this regime, there are no jumps in the PPC (Fig. S2 A), and the activity bump corresponds to the stimuli presented in the previous trial in a fewer fraction of the trials (Fig. S2 C), relative to when adaptation is present (Fig. 3 B). As a result, no short-term history effects can be observed (Fig. S2 C and D), even though a strong contraction bias persists (Fig. S2 E).

**Figure 4:**
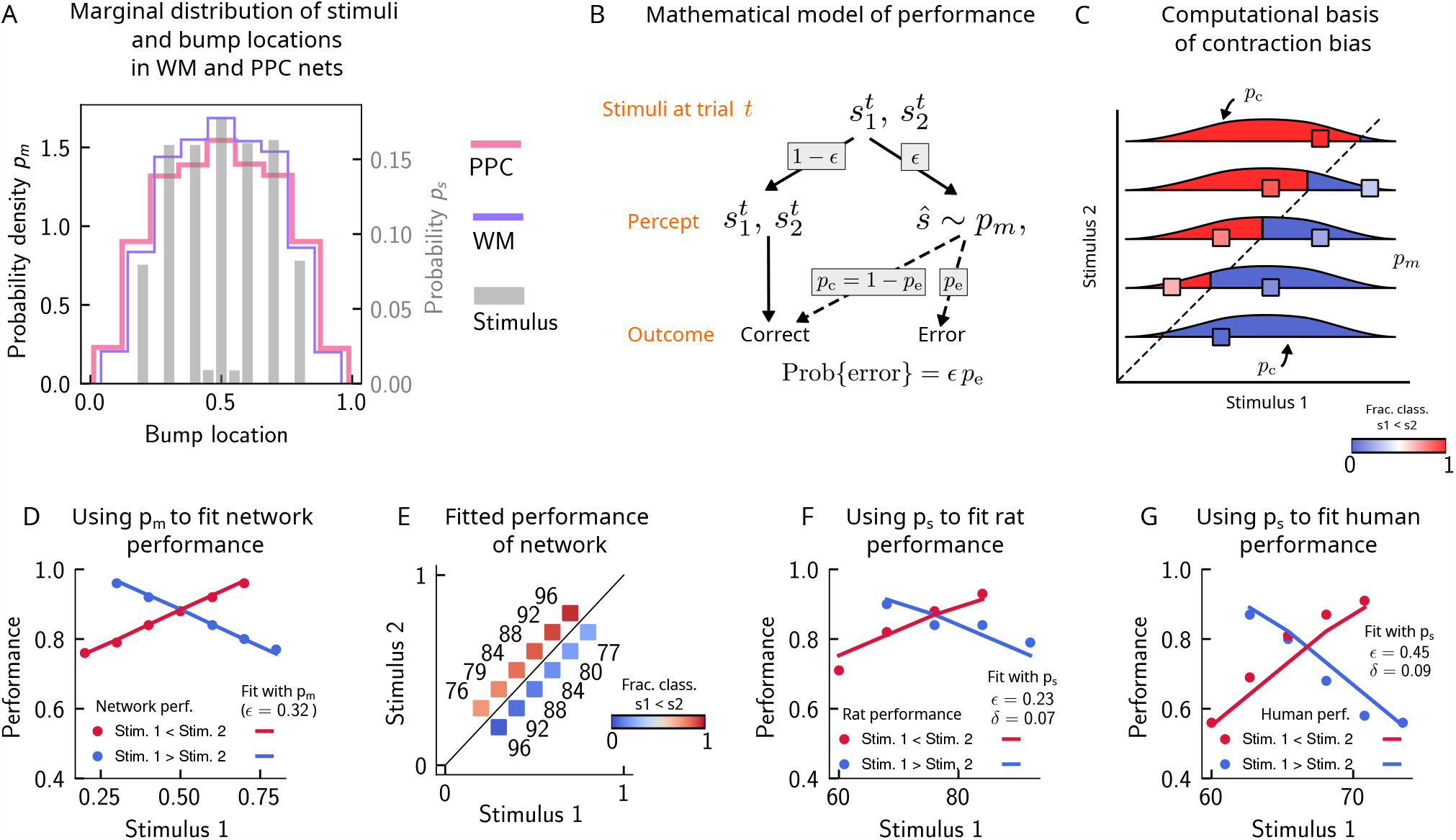
Errors are drawn from the marginal distribution of stimuli, giving rise to contraction bias. **(A)** The bump locations in both the WM network (in pink) and the PPC network (in purple) have identical distributions to that of the input stimulus (marginal over *s*_1_ or *s*_2_, shown in gray). **(B)** A simple mathematical model illustrates how contraction bias emerges as a result of a volatile working memory for *s*_1_. A given trial consists of two stimuli 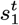 and 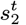 . We assume that the encoding of the second stimulus 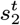 is error-free, contrary to the first stimulus that is prone to change, with probability *ϵ*. Furthermore, when *s*_1_ does change, it is replaced by another stimulus, *ŝ* (imposed by the input from the PPC in our network model). Therefore, *ŝ* is drawn from the marginal distribution of bump locations in the PPC, which is similar to the marginal stimulus distribution (see panel B), *p*_*m*_ (see also Sect. 4.2). Depending on the new location of *ŝ*, the comparison to *s*_2_ can either lead to an erroneous choice (Bias-, with probability *p*_*e*_) or a correct one (Bias+, with probability *p*_*c*_ = 1− *p*_*e*_). **(C)** The distribution of bump locations in PPC (from which replacements *ŝ* are sampled) is overlaid on the stimulus set, and repeated for each value of *s*_2_. For pairs below the diagonal, where *s*_1_ *> s*_2_ (blue squares), the trial outcome will be an error if the displaced WM bump *ŝ* ends up above the diagonal (red section of the *p*_*m*_ distribution). The probability to make an error, *p*_*e*_, equals the integral of *p*_*m*_ over values above the diagonal (red part), which increases as *s*_1_ increases. Vice versa, for pairs above the diagonal (*s*_1_ *< s*_2_, red squares), *p*_*e*_ equals the integral of *p*_*m*_ over values below the diagonal, which increases as *s*_1_ decreases. **(D)** The performance of the attractor network as a function of the first stimulus *s*_1_, in red dots for pairs of stimuli where *s*_1_ *> s*_2_, and in blue dots for pairs of stimuli where *s*_1_ *< s*_2_. The solid lines are fits of the performance of the network using Eq. 9, with *ϵ* as a free parameter. **(E)** Numbers correspond to the performance, same as in (D), while colors expresses the fraction classified as *s*_1_ *< s*_2_ (colorbar), to illustrate the contraction bias. **(F)** Performance of rats performing the auditory delayed-comparison task in Ref.[7]. Dots correspond to the empirical data, while the lines are fits with the statistical model, using the distribution of stimuli. The additional parameter *δ* captures the lapse rate. **(G)** Same as (F), but with humans performing the task. Data in (F) and (G) reproduced with permission from Ref.[7].

As in the study in Ref.[7], we can further study the impact of the PPC on the dynamics of the WM network by weakening the weights from the PPC to the WM network, mimicking the inactivation of PPC (Fig. 2 E and F, Fig. S3 A and B). Under this manipulation, the trajectory of the activity bump in the WM network immediately before the onset of the second stimulus *s*_2_ closely follows the external input, consistent with an enhanced WM function (Fig. S3 C and D).

The drift-jump dynamics in our model of the PPC give rise to short-term (notably one and two-trial back) sensory history effects in the performance of the WM network. In addition, we observe an equally salient contraction bias (bias towards the sensory mean) in the WM network’s performance, increasing with the delay period (Fig. 3 E). However, we find that the activity bump in both the WM and the PPC network corresponds to the mean over all stimuli in only a small fraction of trials, expected by chance (Fig. 3 B, see Sect. 4.1 for how it is calculated). Rather, the bump is located more often at the current trial stimulus 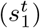, and to a lesser extent, at the location of stimuli presented at the previous trial 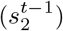. As a result, contraction bias in our model cannot be attributed to the representation of the running sensory average in the PPC. In the next section, we show how contraction bias arises as an averaged effect, when single trial errors occur due to short-term sensory history biases.

### 1.4 Errors are drawn from the marginal distribution of stimuli, giving rise to contraction bias

In order to illustrate the statistical origin of contraction bias in our network model, we consider a mathematical scheme of its performance (Fig. 4 B). In this simple formulation, we assume that the first stimulus to be kept in working memory, 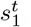, is volatile. As a result, in a fraction *ϵ* of the trials, it is susceptible to replacement with another stimulus *ŝ* (by the input from the PPC, that has a given distribution *p*_*m*_ (Fig. 4 A)). However, this replacement does not always lead to an error, as evidenced by Bias- and Bias+ trials (i.e. those trials in which the performance is affected, negatively and positively, respectively (Fig. 2 B)). For each stimulus pair, the probability to make an error, *p*_*e*_, is integral of *p*_*m*_ over values lying on the wrong side of the *s*_1_ = *s*_2_ diagonal (Fig. 4 C). For instance, for stimulus pairs below the diagonal (Fig. 4 C, blue squares) the trial outcome is erroneous only if *ŝ* is displaced above the diagonal (red part of the distribution). As one can see, the area above the diagonal increases as *s*_1_ increases, giving rise to a gradual increase in error rates (Fig. 4 C). This mathematical model can capture the performance of the attractor network model, as can be seen through the fit of the network performance, when using the bump distribution in the PPC as *p*_*m*_, and *ϵ* as a free parameter (see Eq. 9 in Sect. 4.2, Fig. 4 D, E).

Can this simple statistical model also capture the behavior of rats and humans (Fig. 2 C)? We carried out the same analysis for rats and humans, by replacing the bump location distribution of PPC with that of the marginal distribution of the stimuli provided in the task, based on the observation that the former is well-approximated by the latter (Fig. 4 A). In this case, we see that the model roughly captures the empirical data (Fig. 4 F and G), with the addition of another parameter *δ* that accounts for the lapse rate. Interestingly, such “lapse” also occurs in the network model (as seen by the small amount of errors for pairs of stimuli where *s*_2_ is smallest and largest (Fig. 4 E). This occurs because of the drift present in the PPC network, that eventually, for long enough delay intervals, causes the bump to arrive at the boundaries of the attractor, which would result in an error.

This simple analysis implies that contraction bias in the WM network in our model is not the result of the representation of the mean stimulus in the PPC, but is an effect that emerges as a result of the PPC network’s sampling dynamics, mostly from recently presented stimuli. Indeed, a “contraction to the mean” hypothesis only provides a general account of which pairs of stimuli should benefit from a better performance and which should suffer, but does not explain the gradual accumulation of errors upon increasing (decreasing) *s*_1_, for pairs below (above) the *s*_1_ = *s*_2_ diagonal [38, 39, 7]. Notably, it cannot explain why the performance in trials with pairs of stimuli where *s*_2_ is most distant from the mean stand to benefit the most from it. All together, our model suggests that contraction bias may be a simple consequence of errors occurring at single trials, driven by inputs from the PPC that follow a distribution similar to that of the external input (Fig. 4 B).

### 1.5 Contraction bias in continuous recall

Can contraction bias also be observed in the activity of the WM network prior to binary decision-making? Many studies have evidenced contraction bias also in delayed estimation (or production) paradigms, where subjects must retain the value of a continuous parameter in WM and reproduce it after a delay [15, 42]. Given that we observe contraction bias in the behavior of the network, we reasoned that this should also be evident prior to binary decision-making. Similar to delayed estimation tasks, we therefore analyzed the position of the bump *ŝ*, at the end of the delay interval, for each value of *s*_1_. Consistent with our reasoning, we observe contraction bias of the value of *ŝ*, as evidenced by the systematic departure of the curve corresponding to the bump location from that of the nominal value of the stimulus (Fig. 5 A). We also find that this contraction bias becomes greater as the delay interval increases (Fig. 5 A, right). We next analyzed the effect of the previous trial on the current trial by computing the displacement of the bump during the WM delay, as a function of the distance between the current trial’s stimulus and the previous trial’s stimulus *s*_1_(*t*) − *s*_2_(*t*− 1) (Fig. 5 B). We found that when this distance is larger, the displacement of the bump during WM is on average also larger (Fig. 5 B). This displacement is also attractive. Breaking down these effects by delay, we find that longer delays lead to greater attraction (Fig. 5 B, right).

**Figure 5:**
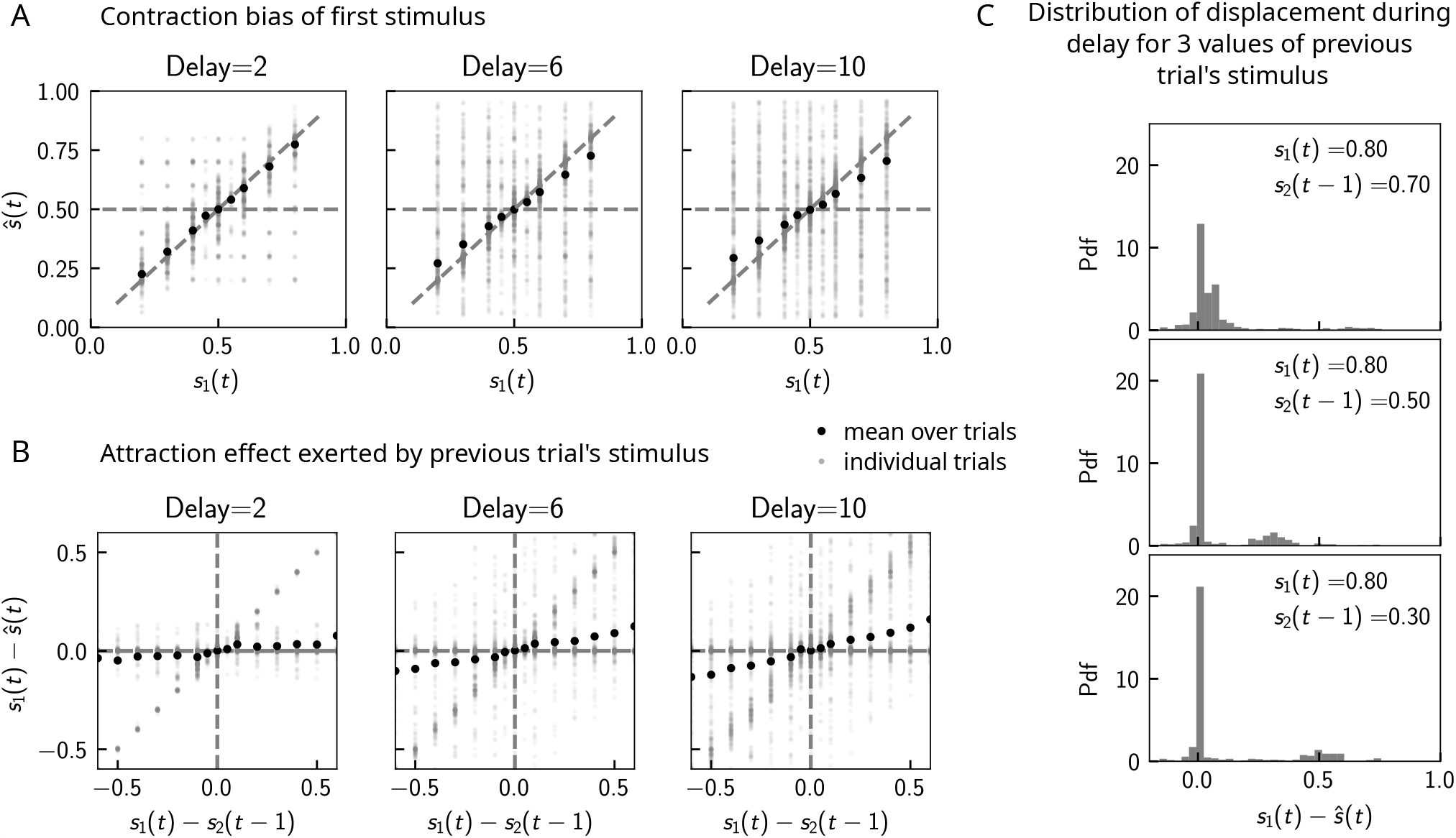
Contraction bias in continuous recall. **(A)** We observe contraction bias of the bump of activity after the delay period ŝ: the average *ŝ* over trials (black dots) deviates from the identity line (diagonal dashed line) toward the mean of the marginal stimulus distribution (0.5). This effect is stronger as the delay interval is longer (left to right panel). **(B)** This contraction bias is actually largely due to the effect of the previous trial: the larger the difference between the current trial and the previous trial’s stimulus *s*_2_(*t−* 1), the larger is this attractive effect on average. Accordingly with panel (A), this effect is stronger for longer delay intervals (left to right panel). **(C)** The distribution of the bump displacement during delay period is characterized by two modes: a main one centered around 0, corresponding to correct trials where the WM bump is not displaced during the delay interval, and another one centered around *s*_1_(*t*) − *s*_2_(*t−* 1), where the bump is displaced during WM (delay interval is randomly selected between 2, 4 and 10 seconds. We show here this distribution for three values of *s*_2_(*t −* 1).

These results point to attractive effects of the previous trial, leading in turn to contraction bias in our model. To better understand the dynamics leading to them, we next looked at the distribution of bump displacements conditioned on a specific value of the second stimulus of the previous trial *s*_2_(*t −*1) (Fig. 5 C). These distributions are characterized by a mode around 0, corresponding to a majority of trials in which the bump is not displaced, and another mode around *s*_1_(*t*) − *s*_2_(*t*−1), corresponding to the displacement in the direction of the preceding trial’s stimulus described in Sect. 1.3 and Fig. 2 B. However, note that the variance of this second mode can be large, reflecting displacements to locations other than *s*_2_(*t−* 1), due to the complex dynamics in both networks that we have described in detail in Sect. 1.3.

### 1.6 Model predictions

#### 1.6.1 The stimulus distribution impacts the pattern of contraction bias through its cumulative

In our model, the pattern of errors is determined by the cumulative distribution of stimuli from the correct decision boundary *s*_1_ = *s*_2_ to the left (right) for pairs of stimuli below (above) the diagonal (Fig. 4 C and Fig. S4 A). This implies that using a stimulus set in which this distribution is deformed makes different predictions for the gradient of performance across different stimulus pairs. A distribution that is symmetric (Fig. S4 A) yields an equal performance for pairs below and above the *s*_1_ = *s*_2_ diagonal (blue and red lines) when *s*_1_ is at the mean (as well as the median, given the symmetry of the distribution). A distribution that is skewed, instead, yields an equal performance when *s*_1_ is at the median for both pairs below and above the diagonal. For a negatively skewed distribution (Fig. S4 B) or positively skewed distribution (Fig. S4 C) the performance curves for pairs of stimuli below and above the diagonal show different concavity. For a distribution that is bimodal, the performance as a function of *s*_1_ resembles a saddle, with equal performance for intermediate values of *s*_1_ (Fig. S4 D). These results indicate that although the performance is quantitatively shaped by the form of the stimulus distribution, it persists as a monotonic function of *s*_1_ under a wide variety of manipulations of the distributions. This is a result of the property of the cumulative function, and may underlie the ubiquity of contraction bias under different experimental conditions.

We compare the predictions from our simple statistical model to the Bayesian model in [41], outlined in Sec. 4.3. We compute the predicted performance of an ideal Bayesian decision maker, using a value of the uncertainty in the representation of the first stimulus (*σ* = 0.12) that yields the best fit with the performance of the statistical model (where the free parameter is *ϵ* = 0.5, Fig. S4 A, B, C, and D, second panels). Our model makes different predictions across all types of distributions from that of the Bayesian model. Across all of the distributions (used as priors, in the Bayesian model), the main difference is that of a monotonic dependence of performance as a function of *s*_1_ for our model (Fig. S4 A, B, C, and D, second panels). The biggest difference can be seen with a prior in which pairs of stimuli with extreme values are much more probable than middle-range values. Indeed, in the case of a bimodal prior, for pairs of stimuli where our model would predict a worse-than-average performance (Fig. S4 D, third panel), the Bayesian model predicts a very good performance (Fig. S4 D, fourth panel).

Do human subjects perform as predicted by our model (Fig. 6 A)? We tested 34 human subjects on the auditory modality of the task. The experimental protocol was identical to the one used in Ref.[7]. Briefly, participants were presented with two sounds separated by a delay interval that varied across trials (randomly selected from 2, 4 and 6 seconds). After the second sound, participants were required to decide which sound was louder by pressing the appropriate key. We tested two groups of participants on two stimulus distributions: a negatively skewed and a bimodal distribution (Fig. 6 A, see Sect. 3.3 for more details). Participants performed the task with a mean accuracy of approximately 75%, across stimulus distribution groups and across pairs of stimuli (Fig. 6 B). The experimental data was compatible with the predictions of our model. First, for the negatively skewed stimulus distribution condition, we observe a shift of the point of equal performance to the right, relative to a symmetric distribution (Fig. 6 C, left panel). For the bimodal condition, such a shift is not observed, as predicted by our model (Fig. 6 C, right panel). Second, the monotonic behavior of the performance, as a function of *s*_1_ also holds, across both distributions (Fig. 6 C). Our model provides a simple explanation: the percent correct on any given pair is given by the probability that, given a shift in the working memory representation, this representation still does not affect the outcome of the trial (Fig. 4 C). This probability, is given by cumulative of the probability distribution of working memory representations, for which we assume the marginal distribution of the stimuli to be a good approximation (Fig. 4 A). As a result, performance is a monotonic function of *s*_1_, independent of the shape of the distribution, while the same does not always hold true for the Bayesian model (Fig. 6 C).

**Figure 6:**
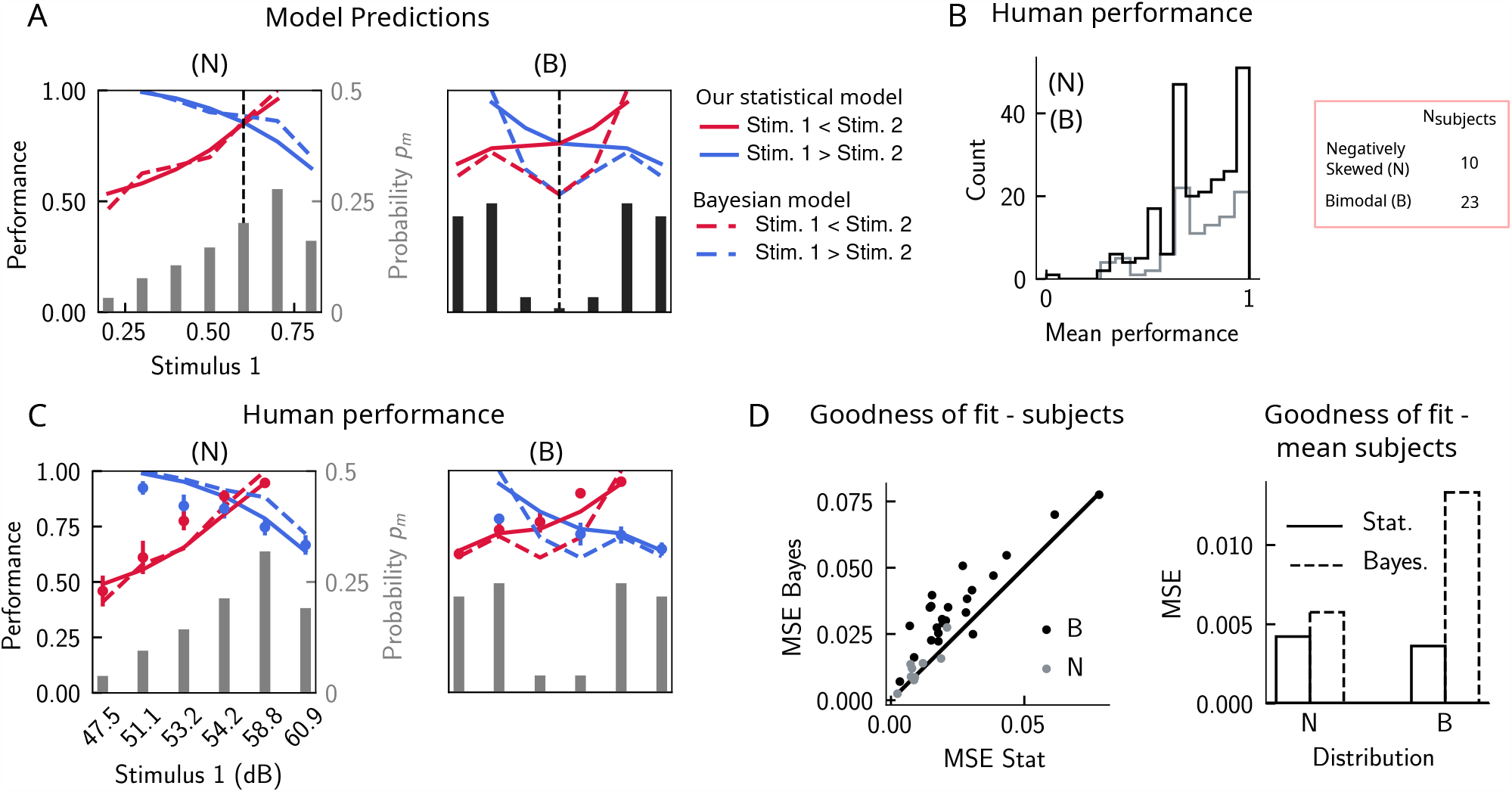
The stimulus distribution impacts the pattern of contraction bias through its cumulative. **(A)** Left panel: prediction of performance (left y-axis) of our statistical model (solid lines) and the Bayesian model (dashed lines) for a negatively skewed stimulus distribution (gray bars, to be read with the right y-axis). Blue (red): performance as a function of *s*_1_ for pairs of stimuli where *s*_1_ *> s*_2_ (*s*_1_ *< s*_2_). Vertical dashed line: median of distribution. Right: same as left, but for a bimodal distribution. **(B)** The distribution of performance across different stimuli pairs and subjects for the negatively skewed (gray) and the bimodal distribution (black). On average, across both distributions, participants performed with an accuracy of 75%. **(C)** Left: mean performance of human subjects on the negatively skewed distribution (dots, error-bars correspond to the standard deviation across different participants). Solid (dashed) lines correspond to fits of the mean performance of subjects with the statistical (Bayesian) model, *ϵ* = 0.55 (*σ* = 0.38). Red (blue): performance as a function of *s*_1_ for pairs of stimuli where *s*_1_ *< s*_2_ (*s*_1_ *> s*_2_), to be read with the left y-axis. The marginal stimulus distribution is shown in gray bars, to be read with the right y-axis. Right: same as left panel, but for the bimodal distribution. Here *ϵ* = 0.54 (*σ* = 0.73). **(D)** Left: goodness of fit, as expressed by the mean-squared-error (MSE) between the empirical curve and the fitted curve (statistical model in the x-axis and the Bayesian model in the y-axis), computed individually for each participant and each distribution. Right: goodness of fit, computed for the average performance over participants in each distribution.

We further fit the performance of each participant, using both our statistical model and the Bayesian model, by minimizing the mean squared error loss (MSE) between the empirical curve and the model, with *ϵ* and *σ* as free parameters (Fig. 6 C), respectively (for the Bayesian model, we used the marginal distribution of the stimuli *p*_*m*_ as the prior). Across participants in both distributions, our statistical model yielded a better fit of the performance, relative to the Bayesian model (Fig. 6 D, left panel). We further fit the mean performance across all participants within a given distribution group, and similarly found that the statistical model yields a better fit, using the MSE as a goodness of fit metric (Fig. 6 D, right panel).

Finally, in order to better understand the parameters that affect the occurrence of errors in human participants, we computed the performance and fraction classified as *s*_1_ *< s*_2_ separately for different delay intervals. We found that the larger the delay interval, the lower the average performance (Fig. 7 A), accompanied by a larger contraction bias for larger intervals (Fig. 7 B). We further analyzed the fraction of trials in which subjects responded *s*_1_ *< s*_2_, conditioned on the specific pair of stimuli presented in the current and the previous trial (Fig. 7 C) for all distributions (one negatively skewed, and two bimodal distributions, of which only one is shown in Fig. 6 C). Compatible with the previous results [7], we found attractive history effects that increased with the delay interval (Fig. 7 D and E).

**Figure 7:**
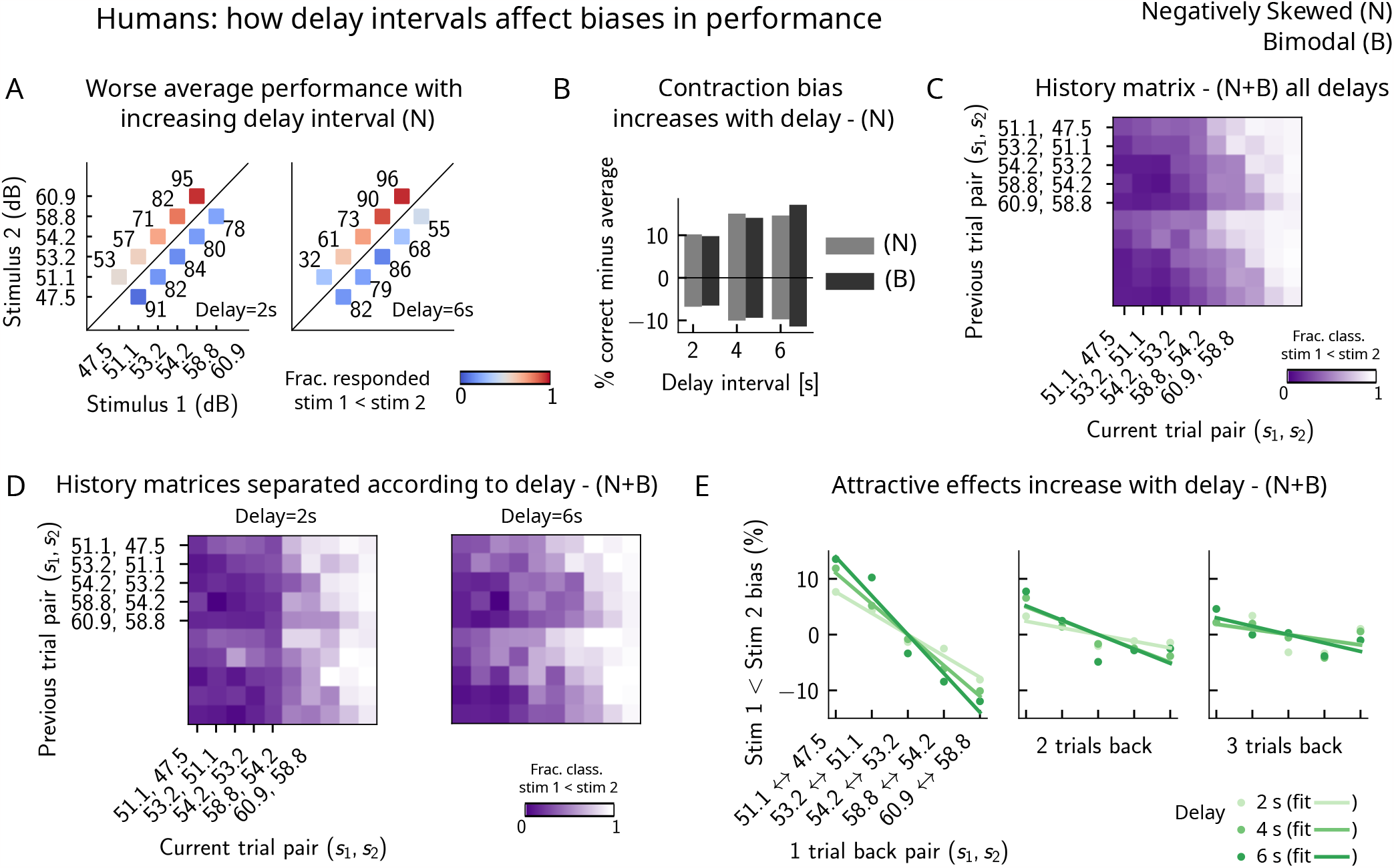
Attractive effects of the previous trials lead to contraction bias in human subjects, both increasing with delay interval. **(A)** The performance (in percentage correct, shown in numbers above each stimulus pair) of human subjects is better with lower delay intervals (left, 2 seconds), than with higher delay intervals (right, 6 seconds). Colorbar expresses the fraction of trials in which participants responded that *s*_1_ *< s*_2_. Results are for the negatively skewed stimulus distribution, noted (N). **(B)** Concurrently, contraction bias on bias+ and bias-trials (quantification explained in text) also increases with an increased delay interval, for both stimulus distributions (negatively skewed in gray and bimodal in black). **(C)** History matrix, expressing the fraction of trials in which subjects responded *s*_1_ *< s*_2_ (in color) for every pair of current (x-xis) and previous (y-axis) stimuli, for negatively skewed and bimodal stimulus distributions (N+B). The one-trial back history effects can be seen through the vertical modulation of the color. Colorbar codes for the fraction of trials in which subjects responded *s*_1_ *< s*_2_. **(D)** History matrices (as in (C)), computed for all distributions, and separated according to delay intervals (left: 2 seconds and right: 6 seconds). **(E)** Bias, quantifying the (attractive) effect of previous stimulus pairs, for 1− 3 trials back in history. The attractive bias, computed for all distributions, increases with the delay interval separating the two stimuli (light to dark green: increasing delay).

#### 1.6.2 A prolonged inter-trial interval improves average performance and reduces attractive bias

If errors are due to the persistence of activity resulting from previous trials, what then, is the effect of the inter-trial interval (ITI)? In our model, a shorter ITI (relative to the default value of 6*s* used in Figs. 2 and 3) results in a worse performance and vice versa (Fig. 8 A, B, C). This change in performance is reflected in reduced biases toward the previous trial (Fig. 8 D and E). A prolonged ITI allows for a drifting bump to vanish due to the effect of adaptation: as a result, the performance improves with increasing ITI and conversely, worsens with a shorter ITI.

**Figure 8:**
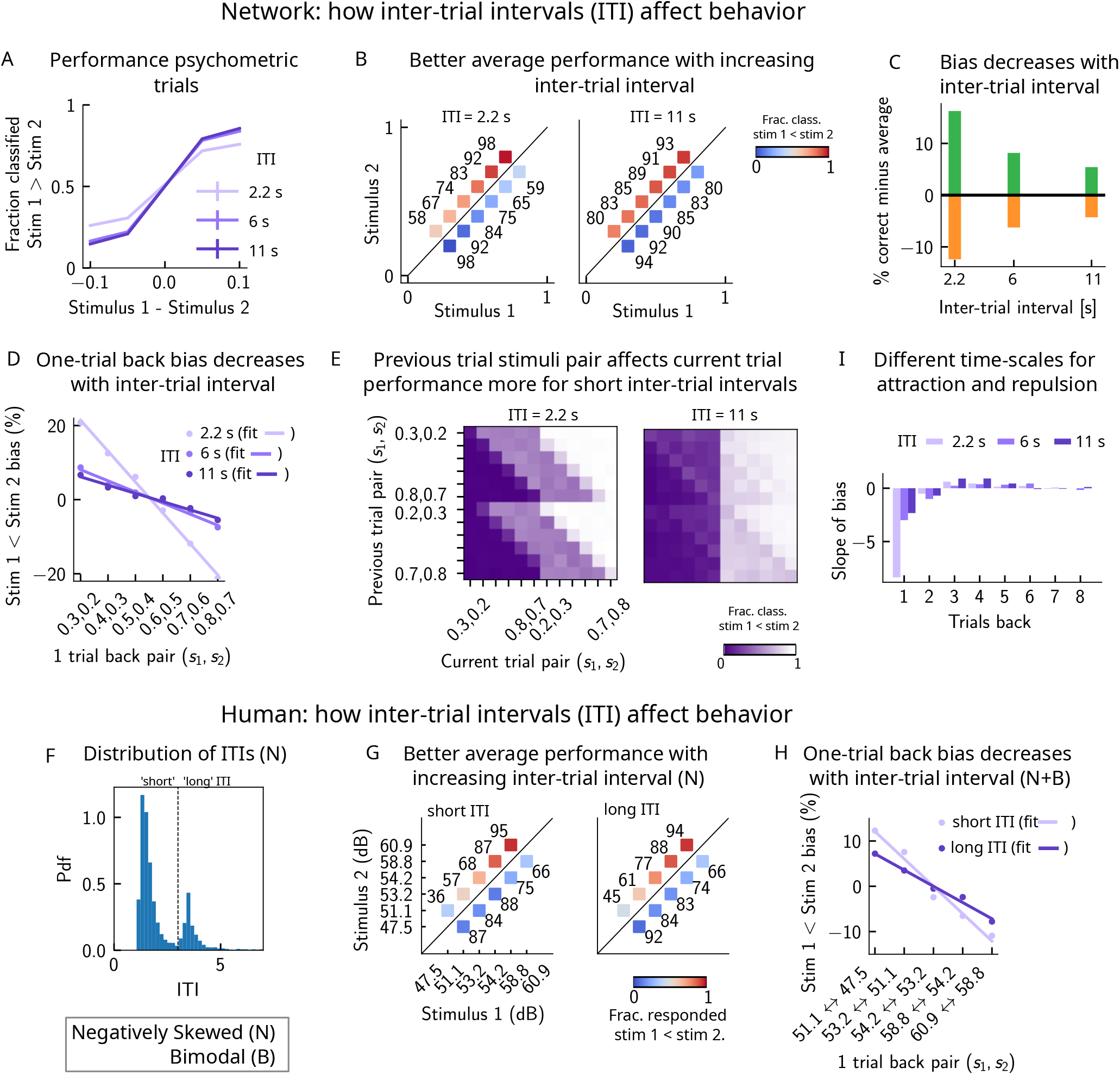
A prolonged inter-trial interval (ITI) improves average performance and reduces attractive biases. Working memory is attracted towards short-term and repelled from long-term sensory history. **(A)** Performance of the network model for the psychometric stimuli improves with an increasing inter-trial interval. Errorbars (not visible) correspond to the s.e.m. over different simulations. **(B)** The network performance (numbers next to stimulus pairs) is on average better for longer ITIs (right panel, ITI=11*s*), compared to shorter ones (left panel, ITI=2.2*s*). Colorbar indicates the fraction of trials classified as *s*_1_ *< s*_2_. **(C)** Quantifying contraction bias separately for bias+ trials (green) and bias-trials (orange) yields a decreasing bias as the inter-trial interval increases. **(D)** The bias, quantifying the (attractive) effect of the previous trial, decreases with ITI. Darker shades of purple correspond to increasing values of the ITI, with dots corresponding to simulation values and lines to linear fits. **(E)** Performance is modulated by the previous stimulus pairs (modulation along the y-axis), more for a short ITI (left, ITI=2.2*s*) than for a longer ITI (right, ITI=11*s*). The colorbar corresponds to the fraction classified *s*_1_ *< s*_2_. **(F)** The distribution of ITIs in the human experiment is bimodal. We define as having a “short” ITI, those trials where the preceding ITI is shorter than 3 seconds and conversely for “long” ITI. **(G)** The human performance for the negatively skewed stimulus distribution is on average worse for shorter ITIs (left panel), compared to longer ones (right panel). Colorbar indicates the fraction of trials subjects responded *s*_1_ *< s*_2_. **(H)** The bias, quantifying the (attractive) effect of the previous trial, increases with ITI in human subjects. Darker shades of purple correspond to increasing values of the ITI, with dots corresponding to empirical values and lines to linear fits. **(I)** Although the stimuli shown up to two trials back yield attractive effects, those further back in history yield repulsive effects, notably when the ITI is larger. Such repulsive effects extend to up to 6 trials back.

Do human subjects express less bias with longer ITIs, as predicted by our model? In our simulations, we set the ITI to either 2.2, 6 or 11 seconds, whereas in the experiment, since it is self-paced, the ITI can vary considerably. In order to emulate the simulation setting as closely as possible, we divided trials into two groups: “short” ITIs (shorter than 3 seconds), and “long” ITIs (longer than 3 seconds). This choice was motivated by the shape of the distribution of ITIs, which is bimodal, with a peak around 1 second, and another after 3 seconds (Fig. 8 F). Given the shape of the ITI distribution, we did not divide the ITIs into smaller intervals as this would result in too little data in some intervals. In line with our model, we found a better average performance with increasing ITI accompanied by decreasing contraction bias (Fig. 8 G). In order to quantify one-trial back effects, we used data pertaining to all of the distributions we tested – the negatively skewed, and also two bimodal distributions (of which only one was shown in this manuscript, in Fig. 6 C). This allowed us to obtain clear one-trial-back attractive biases, decreasing with increasing ITI (Fig. 8 H), in line with our model predictions (Fig. 8 B and D).

#### 1.6.3 Working memory is attracted towards short-term and repelled from long-term sensory history

Although contraction bias is robustly found in different contexts, surprisingly similar tasks, such as perceptual estimation tasks, sometimes highlight opposite effects, i.e. repulsive effects [43–45]. Interestingly, recent studies have found both effects in the same experiment: in a study of visual orientation estimation [18], it has been found that attraction and repulsion have different timescales; while perceptual decisions about orientation are attracted towards recently perceived stimuli (timescale of a few seconds), they are repelled from stimuli that are shown further back in time (timescale of a few minutes). Moreover, in the same study, they find that the long-term repulsive bias is spatially specific, in line with sensory adaptation [46–48] and in contrast to short-term attractive serial dependence [18]. Given that adaptation is a main feature of our model of the PPC, we sought to determine whether such repulsive effects can emerge from the model. We extended the calculation of the bias to up to ten trials back, and quantified the slope of the bias as a function of the previous trial stimulus pair. We observe robust repulsive effects appear after the third trial back in history, and up to six trials back (Fig. 8 I). In our model, both short-term attractive effects and longer-term repulsive effects can be attributed to the multiple timescales over which the networks operate. The short-term attractive effects occur due to the long time it takes for the adaptive threshold to build up in the PPC, and the short timescale with which the WM network integrates input from the PPC. The longer-term repulsive effects occur when the activity bump in the PPC persists in one location and causes adaptation to slowly build up, effectively increasing the activation threshold. The raised threshold takes equally long to return to baseline, preventing activity bumps to form in that location and thereby creating repulsion toward all the other locations in the network. Crucially, however, the amplitude of such effects depend on the inter-trial interval; in particular, for shorter inter-trial intervals, the repulsive effects are less observable.

### 1.7 The timescale of adaptation in the PPC network can control perceptual biases similar to those observed in dyslexia and autism

In a recent study [16], a similar PWM task with auditory stimuli was studied in human neurotypical (NT), autistic spectrum (ASD) and dyslexic (DYS) subjects. Based on an analysis using a generalized linear model (GLM), a double dissociation between different subject groups was suggested: ASD subjects exhibit a stronger bias towards long-term statistics – compared to NT subjects –, while for DYS subjects, a higher bias is present towards short-term statistics.

We investigated our model to see if it is able to show similar phenomenology, and if so, what are the relevant parameters controlling the timescale of the biases in behavior. We identified the adaptation timescale in the PPC as the parameter that affects the extent of the short-term bias, consistent with previous literature [49], [50]. Calculating the mean bias towards the previous trial stimulus pair (Fig. 9 A), we find that a shorter-than-NT adaptation timescale yields a larger bias towards the previous trial stimulus. Indeed, a shorter timescale for neuronal adaptation implies a faster process for the extinction of the bump in PPC – and the formation of a new bump that remains stable for a few trials – producing “jumpier” dynamics that lead to a larger number of one-trial back errors. In contrast, increasing this timescale with respect to NT gives rise to a stable bump for a longer time, ultimately yielding a smaller short-term bias. This can be seen in the detailed breakdown of the network’s behavior on the current trial, when conditioned on the stimuli presented at the previous trial (Fig. 9 B, see also Sect. 1.3 for a more detailed explanation of the dynamics). We performed a GLM analysis as in Ref. [16] to the network behavior, with stimuli from four trials back and the mean stimulus as regressors (see Sect. 4.4). This analysis shows that a reduction in the PPC adaptation timescale with respect to NT, produces behavioral changes qualitatively compatible with data from DYS subjects; on the contrary, an increase of this timescale yields results consistent with ASD data (Fig. 9 C).

**Figure 9:**
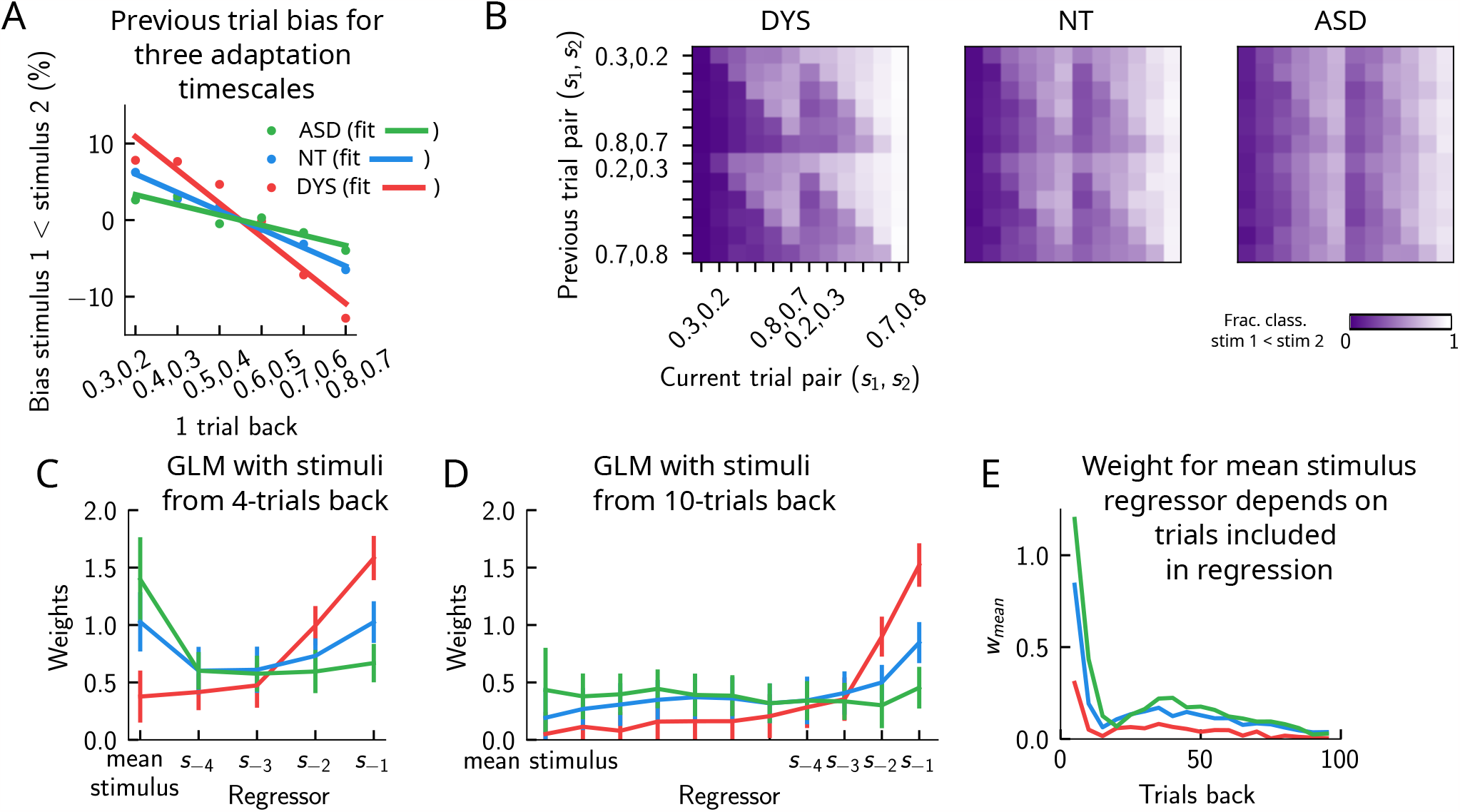
Apparent trade-off between short- and long-term biases, controlled by the timescale of neural adaptation. **(A)** the bias exerted on the current trial by the previous trial (see main text for how it is computed), for three values of the adaptation timescale that mimic similar behavior to the three cohorts of subjects. **(B)** As in (Fig. 2 D), for three different values of adaptation timescale. The colorbar corresponds to the fraction classified *s*_1_ *< s*_2_. **(C)** GLM weights corresponding to the three values of the adaptation parameter marked in (Fig. S6 A), including up to 4 trials back. In a GLM variant incorporating a small number of past trials as regressors, the model yields a high weight for the running mean stimulus regressor. Errorbars correspond to the standard deviation across different simulations. **(D)** Same as in C, but including regressors corresponding to the past 10 trials as well as the running mean stimulus. With a larger number of regressors extending into the past, the model yields a small weight for the running mean stimulus regressor. Errorbars correspond to the standard deviation across different simulations. **(E)** The weight of the running mean stimulus regressor as a function of extending the number of past trial regressors decays upon increasing the number of previous-trial stimulus regressors.

This GLM analysis suggests that dissociable short- and long-term biases may be present in the network behavior. Having access to the full dynamics of the network, we sought to determine how it translates into such dissociable short- and long-term biases. Given that all the behavior arises from the location of the bump on the attractor, we quantified the fraction of trials in which the bump in the WM network, before the onset of the second stimulus, was present in the vicinity of any of the previous trial’s stimuli (Fig. S6 B, right panel, and C), as well as the vicinity of the mean over the sensory history (Fig. S6 B, left panel, and C). While the bump location correlated well with the GLM weights corresponding to the previous trial’s stimuli regressor (comparing the right panels of Fig. S6 A and B), surprisingly, it did not correlate with the GLM weights corresponding to the mean stimulus regressor (comparing the left panels of Fig. S6 A and B). In fact, we found that the bump was in a location given by the stimuli of the past two trials, as well as the mean over the stimulus history, in a smaller fraction of trials, as the adaptation timescale parameter was made larger (Fig. S6 C).

Given that the weights, after four trials in the past, were still non-zero, we extended the GLM regression by including a larger number of past stimuli as regressors. We found that doing this greatly reduced the weight of the mean stimulus regressor (Fig. 9 C, D and E, see Sect. 4.4 and 4.4 for more details). Therefore, we propose an alternative interpretation of the GLM results given in Ref. [16]. In our model, the increased (reduced) weight for long-term mean in the ASD (DYS) subjects can be explained as an effect of a larger (smaller) window in time of short-term biases, without invoking a double dissociation mechanism (Fig. 9 D and E). In Sect. 4.4, we provide a mathematical argument for this, which is empirically shown by including a large number of individual stimuli from previous trials in the regression analysis.

## 2 Discussion

### 2.1 Contraction bias in the delayed comparison task: simply a statistical effect or more?

Contraction bias is an effect emerging in working memory tasks, where in the averaged behavior of a subject, the magnitude of the item held in memory appears to be larger than it actually is when it is “small” and, vice-versa, it appears to be smaller when it is “large” [51, 3, 4, 52, 19, 53, 54]. Recently, Akrami et al [7] have found that contraction bias as well as short-term history-dependent effects occur in an auditory delayed comparison task in rats and humans: the comparison performance in a given trial, depends on the stimuli shown in preceding trials (up to three trials back) [7], similar to previous findings in human 2AFC paradigms [5]. These findings raise the question: does contraction bias occur independently of short-term history effects, or does it emerge as a result of the latter?

Akrami et al [7] have also found the PPC to be a critical node for the generation of such effects, as its optogenetic inactivation (specifically during the delay interval) greatly attenuated both effects. WM was found to remain intact, suggesting that its content was perhaps read-out in another region. Electrophysiological recordings as well as optogenetic inactivation results in the same study suggest that while sensory history information is provided by the PPC, its integration with the WM content must happen somewhere downstream to the PPC. Different brain areas can fit the profile. For instance there are known projections from the PPC to mPFC in rats [55], where neural correlates of parametric working memory have been found [40]. Building on these findings, we suggest a minimal two-module model aimed at better understanding the interaction between contraction bias and short-term history effects. These two modules capture properties of the PPC (in providing sensory history signals) and a downstream network holding working memory content. Our WM and PPC networks, despite having different timescales, are both shown to encode information about the marginal distribution of the stimuli (Fig. 4 A). Although they have similar activity distributions to that of the external stimuli, they have different memory properties, due to the different timescales with which they process incoming stimuli. The putative WM network, from which information to solve the task is read-out, receives additional input from the PPC network. The PPC is modelled as integrating inputs slower relative to the WM network, and is also endowed with firing rate adaptation, the dynamics of which yield short-term history biases and consequently, contraction bias.

It must be noted, however, that short-term history effects (due to firing rate adaptation) do not necessarily need to be invoked in order to recover contraction bias: as long as errors are made following random samples from a distribution in the same range as that of the stimuli, contraction bias should be observed [56]. Indeed, when we manipulated the parameters of the PPC network in such a way that short-term history effects were eliminated (by removing the firing-rate adaptation), contraction bias persisted. As a result, our model suggests that contraction bias may not simply be given by a regression towards the mean of the stimuli during the inter-stimulus interval [57, 58], but brought about by a richer dynamics occurring at the level of individual trials [2], more in line with the idea of random sampling [59].

The model makes predictions as to how the pattern of errors may change when the distribution of stimuli is manipulated, either at the level of the presented stimuli or through the network dynamics. When we tested these predictions experimentally, by manipulating the skewness of the stimulus distribution, such that the median and the mean were dissociated (Fig. 6 A) the results from our human psychophysics experiments were in agreement with the model predictions. In further support of this, in a recent tactile categorization study [45], where rats were trained to categorize tactile stimuli according to a boundary set by the experimenter, the authors have shown that rats set their decision boundary according to the statistical structure of the stimulus set to which they are exposed. More studies are needed to fully verify the extent to which the statistical structure of the stimuli affect the performance. Finally, we note that in our model, the stimulus distribution is not explicitly learned (but see [60]): instead, the PPC dynamics follows the input, and its marginal distribution of activity is similar to that of the external input. This is in agreement with Ref. [45], where the authors used different stimulus ranges across different sessions and noted that rats initiated each session without any residual influence of the previous session’s range/boundary on the current session, ruling out long-term learning of the input structure.

Importantly, our results are not limited to the delayed “comparison” paradigm, where binary decision making occurs. We show that by analyzing the location of the WM bump at the end of the delay interval, similar to the continuous recall tasks, we can retrieve the averaged effects of contraction bias, similar to previous reports [42]. Such continuous readout of the memory reveals a rich dynamics of errors at the level of individual trials, similar to the delayed comparison case, but to our knowledge this has not been studied in previous experimental studies. Papadimitriou et al [15] have characterised residual error distribution, in an orientation recall task, when limiting previous trials to orientations in the range of +35 to +85 degrees relative to the current trial. This distribution is unimodal, leading the authors to conclude that the current trial shows a small but systematic bias toward the location of the memorandum of the previous trial. It remains to be tested whether the error distribution remains unimodal, if conditioned on other values of the current and previous orientations, similar to our analysis in Fig. 5 C.

### 2.2 Attractor mechanism riding on multiple timescales

Our model assumes that the stimulus is held in working memory through the persistent activity of neurons, building on the discovery of persistent selective activity in a number of cortical areas, including the prefrontal cortex (PFC), during the delay interval [61–67]. To explain this finding, we have used the attractor framework, in which recurrently connected neurons mutually excite one another to form reverberation of activity within populations of neurons coding for a given stimulus [68–70]. However, subsequent work has shown that persistent activity related to the stimulus is not always present during the delay period and that the activity of neurons displays far more heterogeneity than previously thought [71]. It has been proposed that short-term synaptic facilitation may dynamically operate to bring a WM network across a phase transition from a silent to a persistently active state [72, 73]. Such mechanisms may further contribute to short-term biases [74], an alternative possibility that we have not specifically considered in this model.

An important model feature that is crucial in giving rise to all of its behavioral effects is its operation over multiple timescales (Fig. S2 F). Such timescales have been found to govern the processing of information in different areas of the cortex [30–32], and may reflect the heterogeneity of connections across different cortical areas [75].

### 2.3 Relation to other models

In many early studies, groups of neurons whose activity correlates monotonically with the stimulus feature, known as “plus” and “minus” neurons, have been found in the PFC [65, 76]. Such neurons have been used as the starting point in the construction of many models [77, 78, 71, 79]. It is important, however, to note that depending on the area, the fraction of such neurons can be small [40], and that the majority of neurons exhibit firing profiles that vary largely during the delay period [80]. Such heterogeneity of the PFC neurons’ temporal firing profiles have prompted the successful construction of models that have not included the basic assumption of plus and minus neurons, but these have largely focused on the plausibility of the dynamics of neurons observed, with little connection to behavior [71].

A separate line of research has addressed behavior, by focusing on normative models to account for contraction bias [19, 5, 59, 81]. The abstract mathematical model that we present (Fig. 4), can be compatible with a Bayesian framework [19] in the limit of a very broad likelihood for the first stimulus and a very narrow one for the second stimulus, and where the prior for the first stimulus is replaced by the distribution of *ŝ*, following the model in Fig. 4 B (see Sect. 4.2 for details). However, it is important to note that our model is conceptually different, i.e. subjects do not have access to the full prior distribution, but only to *samples* of the prior. We show that having full knowledge of the underlying sensory distribution is not needed to present contraction bias effects. Instead, a point estimate of past events that is updated trial to trial suffices to show similar results. This suggests a possible mechanism for the brain to approximate Bayesian inference and it remains open whether similar mechanisms (based on interaction of networks with different integration timescales) can approximate other Bayesian computations. It is also important to note the differences between the predictions from the two models. As shown in Figs. 6 A and S4, depending on the specific sensory distributions, the two models can have qualitatively different testable predictions. Data from our human psychophysical experiments, utilizing auditory Parametric Working Memory, show better agreement with our model predictions as compared to the Bayesian model.

Moreover, an ideal Bayesian observer model alone cannot capture the temporal pattern of short-term attraction and long-term repulsion observed in some tasks, and the model has had to be supplemented with efficient encoding and Bayesian decoding of information in order to capture both effects [18]. In our model, both effects emerge naturally as a result of neuronal adaptation, but their amplitudes crucially depend on the time parameters of the task, perhaps explaining the sometimes contradictory effects reported across different tasks.

Finally, while such attractive and repulsive effects in performance may be suboptimal in the context of a task designed in a laboratory setting, this may not be the case in more natural environments. For example, it has been suggested that integrating information over time serves to preserve perceptual continuity in the presence of noisy and discontinuous inputs [6]. This continuity of perception may be necessary to solve more complex tasks or make decisions, particularly in a non-stationary environment, or in a noisy environment.

## 3 Methods

### 3.1 The model

Our model is composed of two populations of *N* neurons, representing the PPC network and the putative WM network. We consider that each population is organized as a continuous line attractor, with recurrent connectivity described by an interaction matrix *J*_*ij*_, whose entries represent the strength of the interaction between neuron *i* and *j*. The activation function of the neurons is a logistic function, i.e. the output *r*_*i*_ of neuron *i*, given the input *h*_*i*_, is

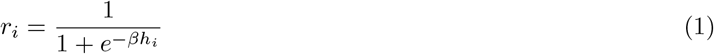

where *β* is the neuronal gain. The variables *r*_*i*_ take continuous values between 0 and 1, and represent the firing rates of the neurons. The input *h*_*i*_ to a neuron is given by

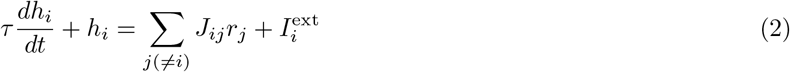

where *τ* is the timescale for the integration of inputs. In the first term on the right hand side, *J*_*ij*_*r*_*j*_ represents the input to neuron *i* from neuron *j*, and 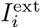 corresponds to the external inputs. The recurrent connections are given by

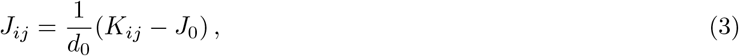

with

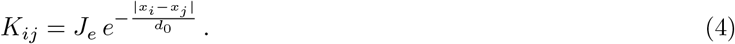

The interaction kernel, *K*, is assumed to be the result of a time-averaged Hebbian plasticity rule: neurons with nearby firing fields will fire concurrently and strengthen their connections, while firing fields far apart will produce weak interactions [82]. Neuron *i* is associated with the firing field *x*_*i*_ = *i/N* . The form of *K* expresses a connectivity between neurons *i* and *j* that is exponentially decreasing with the distance between their respective firing fields, proportional to |*i* − *j*|; the exponential rate of decrease is set by the constant d_0_, i.e. the typical range of interaction. The amplitude of the kernel is also rescaled by *d*_0_, in such a way that ∑ _*i,j*_ *K*_*ij*_ is constant. The strength of the excitatory weights is set by *J*_*e*_; the normalization of *K*, together with the sigmoid activation function saturating to 1, implies that *J*_*e*_ is also the maximum possible input received by any neuron due to the recurrent connections. The constant *J*_0_, instead, contributes to a linear global inhibition term. Its value needs to be chosen depending on *J*_*e*_ and *d*_0_, so that the balance between excitatory and inhibitory inputs ensures that the activity remains localized along the attractor, i.e. it does not either vanish or equal 1 everywhere; together, these three constants set the width of the bump of activity.

The two networks in our model are coupled through excitatory connections from the PPC to the WM network. Therefore, we introduce two equations analogous to Eq. (2), one for each network. The coupling between the two will enter as a firing-rate dependent input, in addition to *I*^ext^. The dynamics of the input to a neuron in the WM network writes

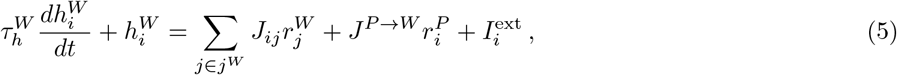

where *j*^*W*^ indexes neurons in the WM network, and 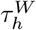 is the timescale for the integration of inputs in the WM network. The first term in the r.h.s corresponds to inputs from recurrent connections within the WM network. The second term, corresponds to inputs from the PPC network. Finally, the last term corresponds to the external inputs used to give stimuli to the network. Similarly, for the PPC network we have

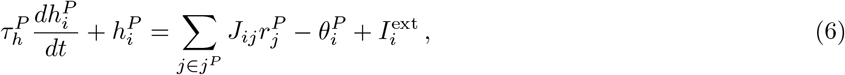

where *j*^*P*^ indexes neurons in the PPC, and where *τ* ^*P*^ is the timescale for the integration of inputs in the PPC network; importantly, we set this to be longer than the analogous quantity for the WM network, 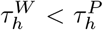 (see Tab. 1). The first and third terms in the r.h.s are analogous to the corresponding ones for the WM network: inputs from within the network and from the stimuli. The second term instead, corresponds to adaptive thresholds with dynamics specified by

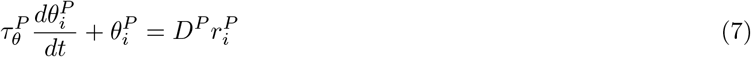

modelling neuronal adaptation, where 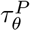 and *D*^*P*^ set its timescale and its amplitude. We are interested in the condition where the timescale of the evolution of the input current is much smaller relative to that of the adaptation 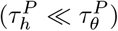. For a constant 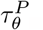, we find that depending on the value of *D*^*P*^, the bump of activity shows different behaviors. For low values of *D*^*P*^, the bump remains relatively stable (Fig. S1 C (1)). Upon increasing *D*^*P*^, the bump gradually starts to drift (Fig. S1 C (2-3)). Upon increasing *D*^*P*^ even further, a phase transition leads to an abrupt dissipation of the bump (Fig. S1 C (4)).

Note that, while the transition from bump stability to drift occurs gradually, the transition from drift to dissipation is abrupt. This abruptness in the transition from the drift to the dissipation regime may imply that only one of the two behaviors is possible in our model of the PPC (Sect. 1.3). In fact, our network model of the PPC operates in the “drift” regime 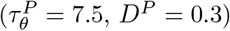. However, we also observe dissipation of the bump, which is mainly responsible for the jumps observed in the model. This occurs due to the inputs from incoming external stimuli, that affect the bump via the global inhibition in the model (Fig. S1 A). Therefore external stimuli can allow the network to temporarily cross the sharp drift/dissipation boundary shown in Fig. S1 B. As a result, the combined effect of adaptation, together with external inputs and global inhibition result in the drift/jump dynamics described in the main text.

Finally, both networks have a linear geometry with free boundary conditions, i.e. no condition is imposed on the profile activity at neuron 1 or *N* .

### 3.2 Simulation

We performed all the simulations using custom Python code. Differential equations were numerically integrated with a time step of *dt* = 0.001 using the forward Euler method. The activity of neurons in both circuits were initialized to *r* = 0. Each stimulus was presented for 400 ms. A stimulus is introduced as a “box” of unit amplitude and of width 2 *δs* around *s* in stimulus space: in a network with *N* neurons, the stimulus is given by setting 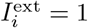 in Eq. 5 for neurons with index *i* within (*s* ± *δs*) × *N*, and 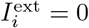 for all the others. Only the activity in the WM network was used to assess performance. To do that, the activity vector was recorded at two time-points: 200 ms before and after the onset of the second stimulus *s*_2_. Then, the neurons with the maximal activity were identified at both time-points, and compared to make a decision. This procedure was done for 50 different simulations with 1000 consecutive trials in each, with a fixed inter-trial interval separating two consecutive trials, fixed to 5 seconds. The inter-stimulus intervals were set according to two different experimental designs, as explained below.

#### 3.2.1 Interleaved design

As in the study in Ref. [7], an inter-stimulus interval of either 2, 6 or 10 seconds was randomly selected. The delay interval is defined as the time elapsed from the end of the first stimulus to the beginning of the second stimulus. This procedure was used to produce Figs. 1, 2, 3, 7, S2, S3.

#### 3.2.2 Block design

In order to provide a comparison to the interleaved design, but also to simulate the design in Ref. [16], we also ran simulations with a block design, where the inter-stimulus intervals were kept fixed throughout the trials. Other than this, the procedure and parameters used were exactly the same as in the interleaved case. This procedure was used to produce Figs. 9 and S6.

### 3.3 Human auditory experiment - delayed comparison task

Subjects received, in each trial, a pair of sounds played from ear-surrounding headphones. The subject self-initiated each trial by pressing the space bar on the keyboard. The first sound was then presented together with a blue square on the left side of a computer monitor in front of the subject. This was followed by a delay period, indicated by ‘WAIT!’ on the screen, then the second sound was presented together with a red square on the right side of the screen. At the end of the second stimulus, subjects had 2 seconds to decide which one was louder, then indicate their choice by pressing the ‘s’ key if they thought that the first sound was louder, or the ‘l’ key if they thought that the second sound was louder. Written feedback about the correctness of their response was provided on the screen for each individual trial. Every ten trials, participants received feedback on their running mean performance calculated up to that trial. Participants then had to press spacebar to go to the next trial (the experiment was hence self-paced).

The two auditory stimuli, *s*_1_ and *s*_2_, separated by a variable delay (of 2, 4 and 6 seconds), were played for 400 ms, with short delay periods of 250 ms inserted before *s*_1_ and after *s*_2_. The stimuli consisted of broadband noise 2000-20000 Hz, generated as a series of sound pressure level (SPL) values sampled from a zero-mean normal distribution. The overall mean intensity of sounds varied from 60-92 dB. Participants had to judge which out of the two stimuli, *s*_1_ and *s*_2_, was louder (had the greater SPL standard deviation).

We recruited 10 subjects for the negatively skewed distribution and 24 subjects for the bimodal distribution. The study was approved by the University College London (UCL) Research Ethics Committee [16159/001] (London, UK). Each participant performed approximately 400 trials for a given distribution. Several participants took part in both distributions.

## 4 Supplementary Material

### 4.1 Computing bump location

In order to check whether the bump is in a target location (Figs. 3 B, S2 B, and S3 D), we check whether the position of the neuron with the maximal firing rate is within a distance of ±5% of the length of the whole line attractor from the target location (Figs. 3 A, S2 A and S3 C). In these figures, we compare the probability that, in a given trial, the activity of the WM network is localized around one of the previous stimuli (estimated from the simulation of the dynamics, histograms) with the probability of this happening due to chance (horizontal dashed line). Here we detail the calculation of the chance probability. In general, if we have two discrete independent random variables, 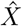 and *Ŷ*, with probability distributions *p*_*X*_ and *p*_*Y*_, the probability of them having the same value is

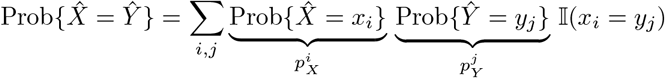

where *i, j* are the indices for different values of the two random variables and 𝕀(*x*_*i*_ = *y*_*j*_) equals 1 where *x*_*i*_ = *x*_*j*_ and 0 otherwise. If the two random variables are identically distributed, the above expression writes

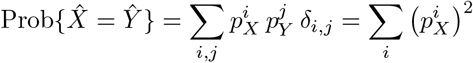

In our case, the two identically distributed random variables are “bump location at the current trial” and the “target bump location” (that are 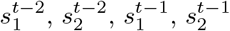 and ⟨*s*⟩). With the exception of the mean stimulus ⟨*s*⟩, all the other variables are identically distributed, with probability *p*_*m*_ (that is the marginal distribution over *s*_1_ or *s*_2_). We note that the bump location in the WM network follows a very similar distribution to *p*_*m*_ (Fig. 4 A). Then, we compute the chance probability with the above relationship, where *p*_*X*_ ≡ *p*_*m*_. For the mean stimulus, instead, we have a probability which is simply equal to 1 for *s* = 0.5 and 0 elsewhere; therefore, the chance probability for the bump location to be at the mean stimulus, then is *p*_*m*_(0.5).

The excess probability (with respect to chance) for the bump location to equal one of the previous stimuli gives a measure of the correlation between these two; in other terms, of the amount of information retained by the network about previous stimuli.

### 4.2 The probability to make errors is proportional to the cumulative distribution of the stimuli, giving rise to contraction bias

In order to illustrate the statistical origin of contraction bias consistent with our network model, we consider a simplified mathematical model of its performance (Fig. 4 B). By definition of the delayed comparison task, the optimal decision maker produces a label *y* equal to 1 if 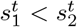, and 0 if 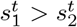; the impossible cases 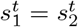 are excluded from the set of stimuli, but would produce a label which is either 0 or 1 with 50% probability. That is

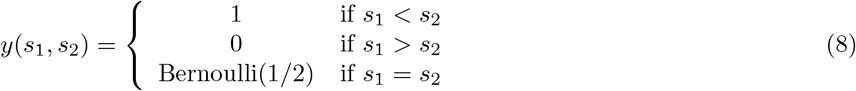

In this simplified scheme, at each trial *t*, the two stimuli 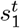 and 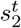 are perfectly perceived with a finite probability 1− *ϵ*, with *ϵ <* 1. Under the assumption that the decision maker behaves optimally based on the perceived stimuli, a correct perception would necessarily lead to the correct label. However, with probability *ϵ*, the first stimulus is randomly selected from a buffer of stimuli, i.e. is replaced by a random variable *ŝ*_1_ that has a probability distribution 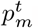.

The probability distribution 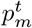 is the statistics of previously shown stimuli. The information about the previous stimulus is given by the activity of the “slower” PPC network. As shown above, after the presentation of the first stimulus of the trial, the bump of activity is seen to jump to the position encoding one of the previously presented stimuli, 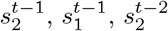 etc. with decreasing probability (Fig. 3 C). Therefore, in calculating the performance in the task, we can take 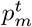 to be the marginal distribution of the stimulus *s*_1_ or *s*_2_ across trials, as in the histogram (Fig. 4 A).

The probability of a misclassification is then given by the probability that, given the pair 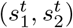, at trial *t*,

1. the first stimulus is replaced by a random value, which happens with probability *ϵ*, and
2. the value of *ŝ*_1_ replaced is larger than 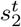 when 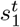 is smaller and viceversa (Fig. 4 C). In summary, the probability of an error at trial *t* is given by

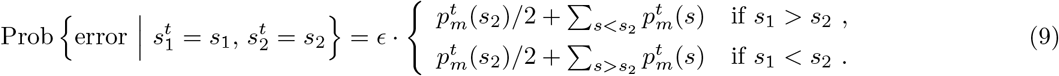

### 4.3 Bayesian description of contraction bias

We reproduce here the theoretical result from [41], which provides a normative model for contraction bias in the Bayesian inference framework, and apply it to the different stimulus distributions described in Sect. 1.6.1.

A stimulus with value *s* is encoded by the agent through a noisy representation 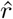 ∼*ℓ*(·|*s*). Before the presentation of the stimulus, the agent has an expectation of its possible values which is described by the probability *π*. Assuming that it has access to the internal representation *r*, as well as the probability distributions *ℓ* and *π*, the agent can infer the perceived stimulusŝ through Bayes rule:

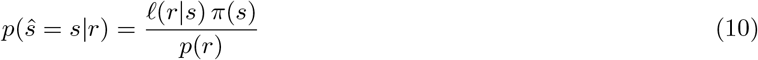

where *p*(*r*) = ∫*ds*^*′*^ *ℓ*(*r|s*^*′*^) *π*(*s*^*′*^). In this Bayesian setting, the probability distributions for the noisy representation and for the expected measurement are interpreted as the *likelihood* and the *prior*, respectively.

In the delayed comparison task, at the time of the decision, the two stimuli *s*_1_ and *s*_2_ are assumed to be encoded independently, although with different uncertainties, due to the different delays leading to the time of decision: *ℓ*(*r*_1_, *r*_2_| *s*_1_, *s*_2_) = *ℓ*_1_(*r*_1_| *s*_1_) *ℓ*_2_(*r*_2|_ *s*_2_), with var[*ℓ*_1_] *>* var[*ℓ*_2_]. Similarly, the expected values of the stimuli are assumed to be independent but also identically distributed: *π*(*s*_1_, *s*_2_) = *π*(*s*_1_) *π*(*s*_2_).

The optimal Bayesian decision maker uses the inference of the stimuli through Eq. (10) to produce an estimate of the probability that *s*_1_ *< s*_2_, given the internal representations,

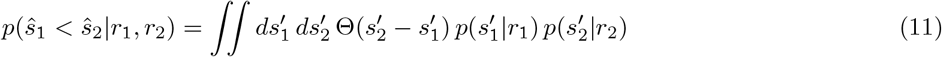

where Θ is the Heaviside function, and yields a label *ŷ*= 1 (truth value of “*s*_1_ *< s*_2_”) when such probability is higher than 1*/*2, and *ŷ* = 0 otherwise. Therefore, the probability that the Bayesian decision maker yields the response “*s*_1_ *< s*_2_” given the *true* values of the stimuli *s*_1_ and *s*_2_ is the average of the label *ŷ* over the possible values of their representations, i.e. over the likelihood:

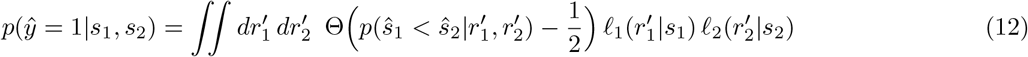

#### 4.3.1 Application to our study

In modelling our data, we assume that the likelihood functions *ℓ*_1_(·| *s*_1_) and *ℓ*_2_(·|*s*_2_) are Gaussian with mean equal to the stimulus, but with different standard deviations, *σ*_1_ and *σ*_2_, respectively, as in [41]. We restrict to the particular case where *σ*_2_ = 0, i.e. there is no uncertainty in the representation of the second stimulus, since there is negligible delay between its presentation and the decision. We instead assume a finite standard deviation *σ*_1_ = *σ*, which we use as the only free parameter of this model to produce Figs.S4 A-D, panels 2 and 4.

The prior *π* is chosen to be the marginal distribution of the first stimulus – identical to the marginal of the second stimulus, because of symmetry.

When *σ*_2_ = 0, *ℓ*_2_(*r*|*s*) = *δ*(*r* − *s*) (Dirac delta), and the predicted response probability, Eq. (12), reduces to

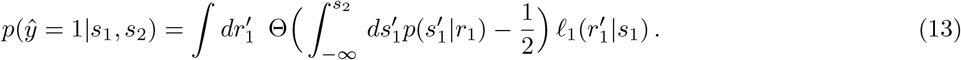

### 4.4 Generalized Linear Model (GLM)

#### GLM as in Lieder et al

Similarly to Ref. [16], we performed a multivariate logistic regression (an instance of generalized linear model, GLM) to the output of the network in the delayed discrimination task with recent stimuli values as covariates:

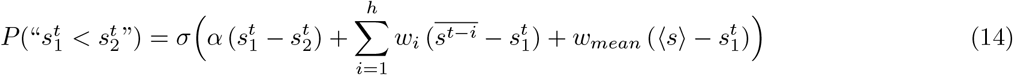

where *σ* is the sigmoidal function 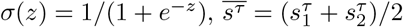 is the mean of the stimuli presented at trial *τ, h* is the number of “history” terms in the regression, and ⟨*s*⟩ is the mean of the stimuli within and across trials up to the current one. As in Ref. [16], we choose *h* = 4, i.e. we include in the short-term history, the four trials prior to the current one. The first term in Eq. (14), with weight *α*, controls the slope of the psychometric curve. The remaining terms, combined linearly with weights *w*, contribute to biases expressing the long and short-term memory. In Ref. [16], it is shown that subjects on the autistic syndrome (ASD) conserve the higher long-term weights, *w*_*mean*_, while losing the short-term weights expressed by neurotypical (NT) subjects. In contrast, dyslexic (DYS) subjects conserve a higher bias from the recent stimuli, *w*_1_, while losing the higher long-term weights, also expressed by neurotypical subjects.

In order to gain insight into this regression model in terms of our network, we also performed a linear regression of the bump of activity just before the onset of the second stimulus, denoted 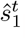, versus the same variables:

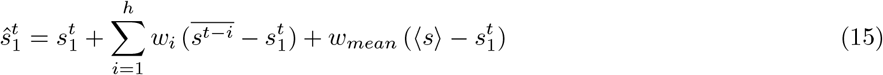

In this case, we see that the weights *w* in the linear regression for 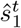 have the same qualitative behavior as the weights for the bias term in the GLM regression for the performance (not shown). This is expected, since the decision-making rule in the network –based on the bump location just before and during the second stimulus, 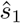 and 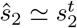 respectively– is deterministic, following 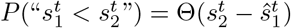. Therefore, the bias term in the GLM performed in Ref. [16], Eq. (14), corresponds to the displacement of the bump location 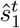 with respect to the actual stimulus 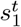, modelled to be linearly dependent on the displacement of previous stimuli from 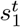.

#### Regression model with infinite history

In the regression formulas in Eqs. (14) and (15), it is possible to give an interpretation of the parameter *w*_*mean*_, that is the weight of the contribution from the covariate corresponding to the mean of the past stimuli. Let us consider two regression models, one in which, in addition to a regressor corresponding to the mean stimulus, regressors corresponding to the stimulus history are included up to trial *h*, and another in which *h* =∞, i.e. infinitely many past stimuli are included as regressors. In this case, Eq. (15) rewrites

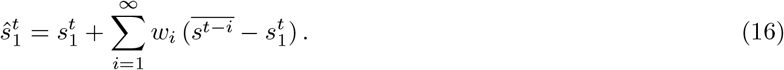

If we assume that the weights obtained from the regression have roughly an exponential dependence on time (Fig. 9 C and D), we can write

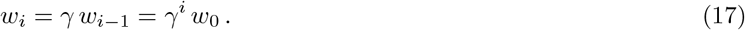

By equating Eqs. (15) and (16), we would find that

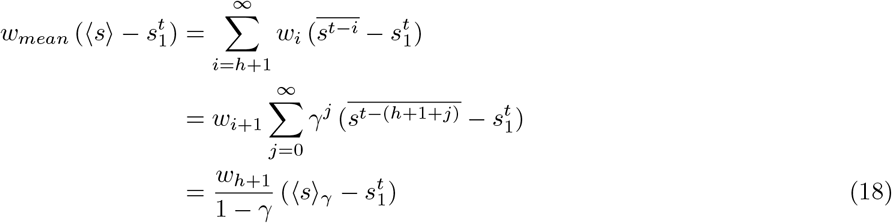

where

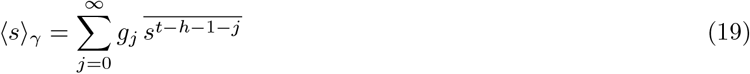

that is an average over the geometric distribution *g*_*j*_ = (1 − *γ*) *γ*^*j*^, from time *t* − (*h* + 1) backward. Since for *γ* large enough we have ⟨*s*⟩_*γ*_ = ⟨*s*⟩, we can identify

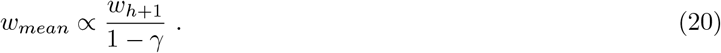

This derivation indicates that the magnitude *w*_*mean*_ in the infinite history model, given by Eq. (15), is a function of the discount factor *γ* as well as the weight of the first trial left out from the finite-history regression (*w*_*h*+1_). A higher *γ* value, i.e. a longer timescale for damping of the weights extending into the stimulus history, yields a higher *w*_*mean*_. We can obtain *γ* for each condition (NT, ASD and DYS) by fitting the weights obtained as a function of trials extending into the history (Fig. 9 C and D). As predicted by Eq. (20), a larger window for short-term history effects (as in the ASD case relative to NT) yields a larger weight for the covariate corresponding to the mean stimulus. Finally, Eq. (20) also predicts that *w*_*mean*_ is proportional to *w*_*h*+1_, the number of trials back we consider in the regression, *h*, implying that the number of covariates that we choose to include in the model may greatly affect the results. Both of these predictions are corroborated by plotting directly the value of *w*_*mean*_ obtained from the regression (Fig. 9 E).

### 4.5 Supplementary Figures

**Figure S1:**
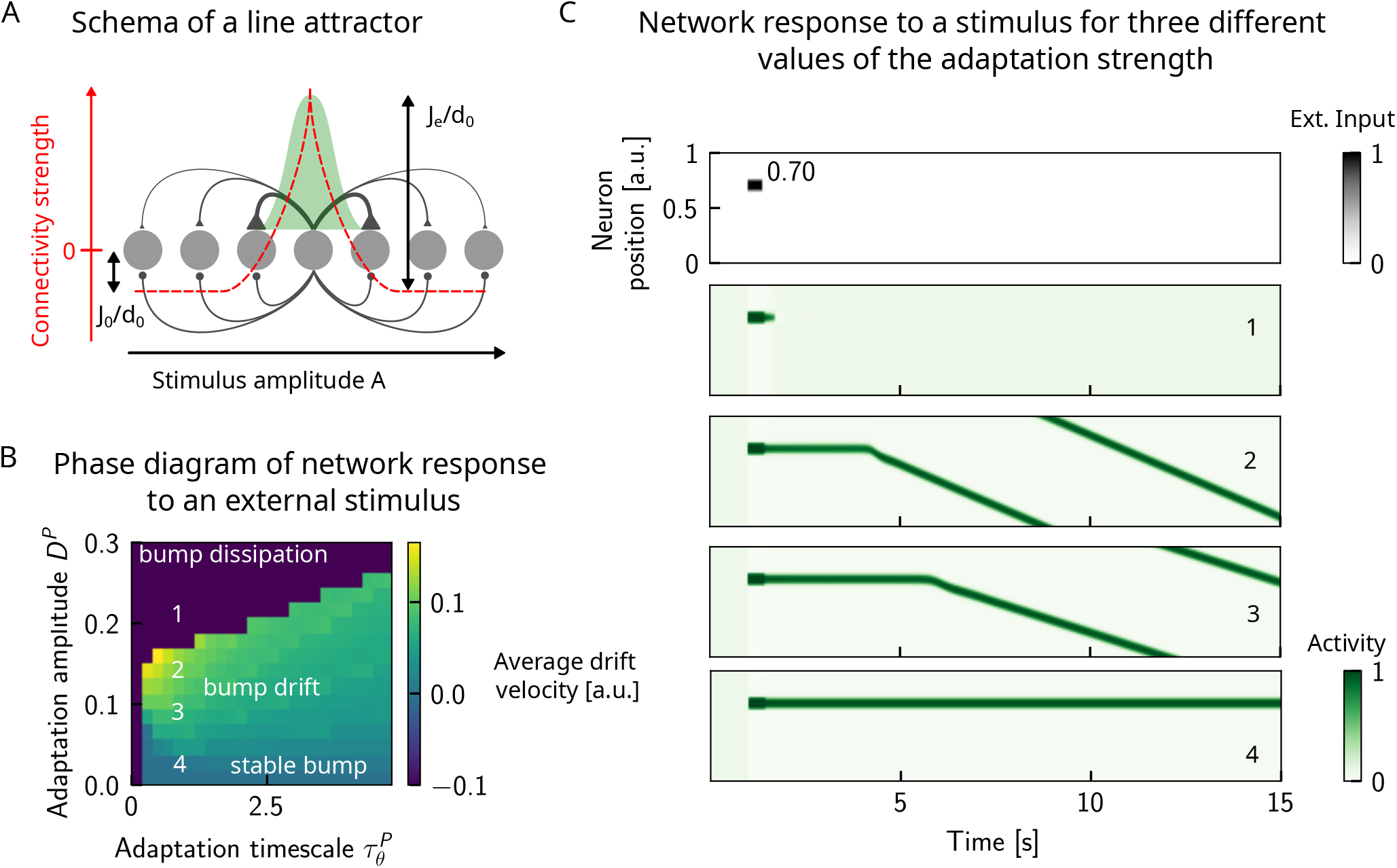
Dynamics of responses in a one-dimensional continuous attractor network, in the presence of adaptation. **(A)** We study a one dimensional line attractor in which neurons code for a stimulus feature that varies along a physical dimension, such as amplitude of an auditory stimulus. The connections between pairs of neurons is a decreasing, symmetric function of the distance between their preferred firing locations, allowing for a bump of activity to form and self-sustain when sufficient input is given to the network. However, this self-sustaining activity may be disrupted if neuronal adaptation is present. In particular, drifting dynamics may be observed. **(B)** Left: phase diagram of the average drift velocity as a function of the adaptation timescale 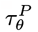 and amplitude *D*^*P*^ . The average drift velocity is simply computed as the distance travelled by the center of the bump in a duration of 50 seconds. Color codes for the average drift velocity (a.u.). Numbers indicate four points for which sample dynamics are shown in (C). **(C)** We observe three main phases: in the first, the activity bump is stable when no or little neuronal adaptation is present (point 4). Larger values of neural adaptation induce drift of the activity bump; the average drift velocity increases upon increasing the neural adaptation (points 2 and 3). Finally, increasing it even further leads to the dissipation of the activity bump (point 1). The boundary between the drift and dissipation phases is abrupt. In these simulations, periodic boundary conditions have been used in order to compute the average average drift velocity over longer durations.

**Figure S2:**
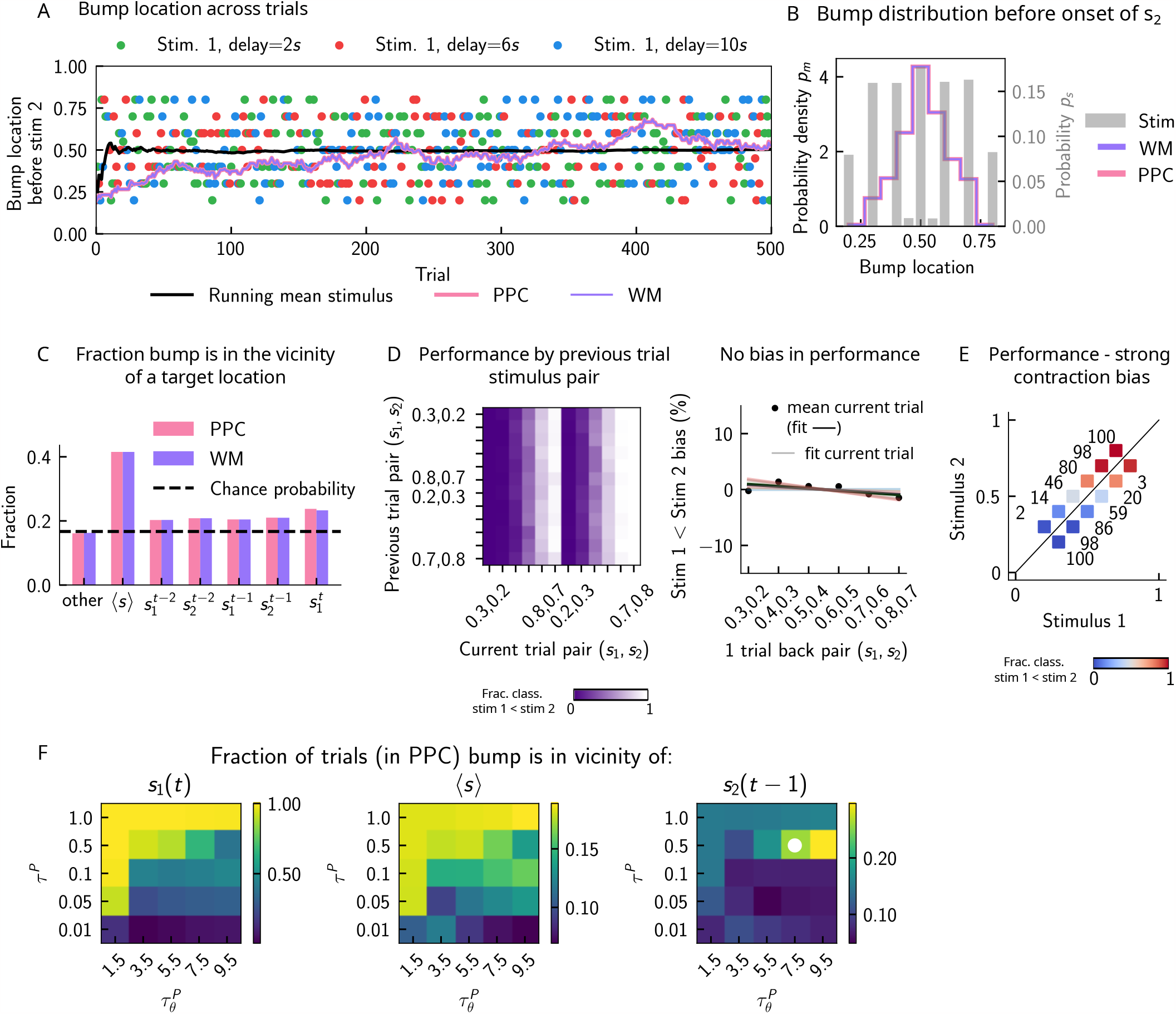
The role of neuronal adaptation in generating short-term history biases. In order to better understand the network mechanisms that give rise to short-term history effects, we removed neural adaptation in the PPC network and assessed the performance in the WM network. **(A)** As in (Fig. 3 B). We track the location of the bump, in the PPC (pink), and in the WM network (purple) before the onset of the second stimulus (the pink curve cannot be seen as the purple curve goes perfectly on top). In this case, the displacement of the bump of activity is smooth and new sensory stimuli (colored dots) induce only a minimal shift in the location of the bump. This behavior is to be contrasted with the case in which there is adaptation in the PPC network, inducing jumps in the bump location (Fig. 3 A). An additional effect of no neural adaptation is that the activity in the PPC network, completely overrides the activity in the WM network. **(B)** As in (Fig. 4 A). Marginal distribution of the bump location in both networks (pink for PPC, purple for WM) before the onset of *s*_2_ is more peaked than the marginal distribution of the stimuli (gray), as a result of the absence of “jumps”. **(C)** As in (Fig. 3 C). We compute the fraction of times the bump is in a given location, current trial 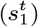, four preceding trials 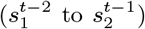, the running mean stimulus, or all other locations (overlapping sets). In this case, in the majority of the trials, the bump is either at the running mean stimulus, or any other location. The fraction of trials in which it is in the position of the four previous stimuli roughly corresponds to chance occurrence (dashed black lines), with only a minor increase for the current stimulus. **(D)** As in (Fig. 2 D). Left: The network behavior conditioned on the previous trial stimulus pair does not exhibit any previous-trial attractive dependence (vertical modulation). Colorbar corresponds to the fraction classified as *s*_1_ *< s*_2_. Right: This attractive dependence can also be expressed through the bias measure (see main text for how it is computed). Colored lines correspond to current trial pairs, the black dots to the mean over all current trial pairs, and the black line to its linear fit. **(E)** As in (Fig. 2 C). Although there are no attractive previous trial effects, the performance expresses a very strong contraction bias, and performance is as if the decision boundary is orthogonal to the optimal decision boundary. Color codes for fraction of trials in which a *s*_1_ *> s*_2_ classification is made. **(F)** Phase diagram with 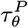 on the x-axis, *τ* ^*P*^ on the y-axis, and in color, the fraction of trials in which the bump, before the network is stimulated with the second stimulus, is in the vicinity of a target (specified in the title of each panel). White dot corresponds to parameters of the default network Tab. 1.

**Figure S3:**
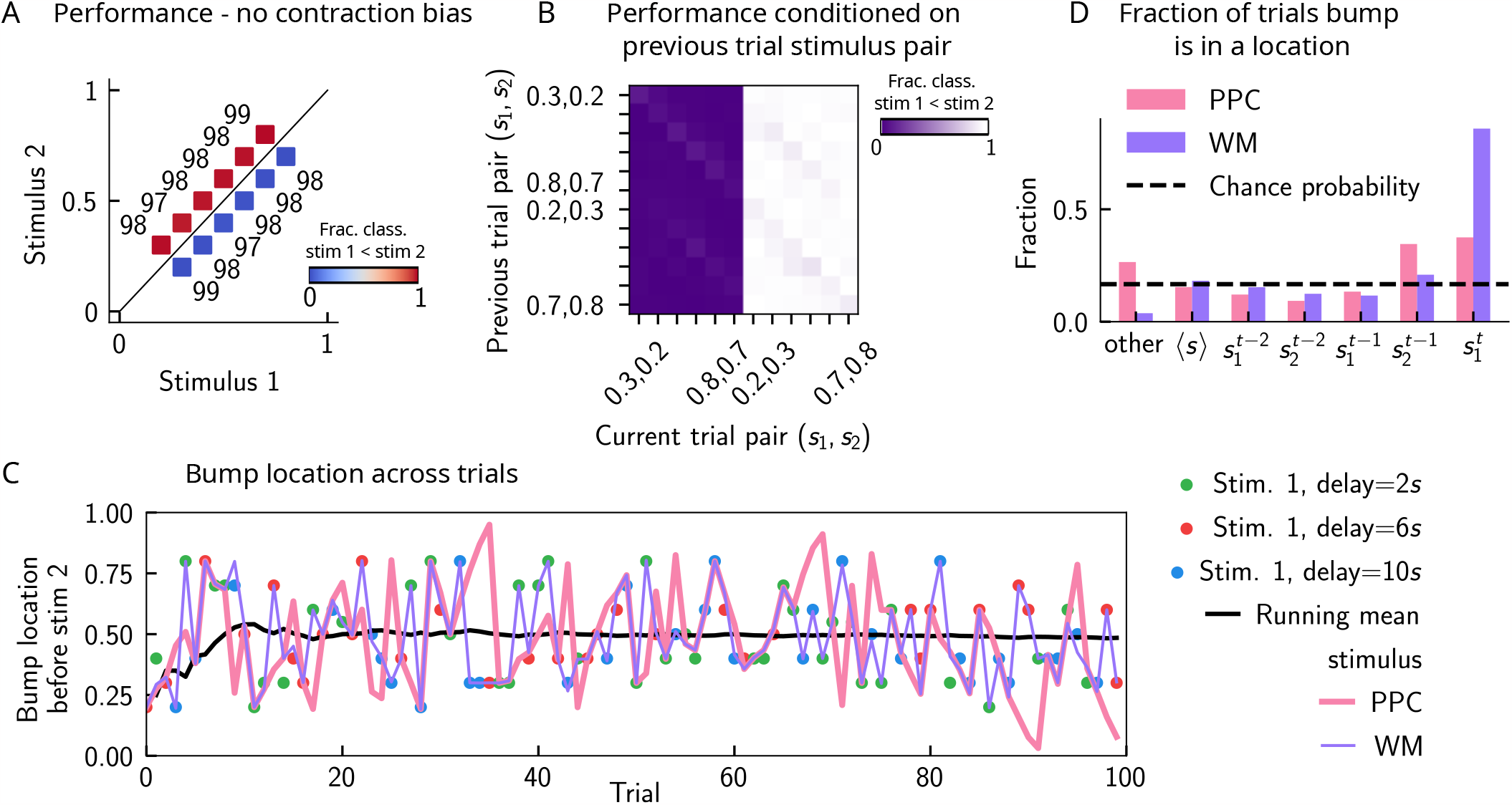
Inactivating the inputs from the PPC network improves performance, in line with experimental findings. **(A)** As in (Fig. 2 C). The performance of the network when the strength of the inputs from the PPC to the WM network is weakened (modelling the optogenetic inactivation of the PPC) is dramatically improved, and contraction bias is virtually eliminated. The colorbar corresponds to the fraction classified as *s*_1_ *< s*_2_. **(B)** As in (Fig. 2 D). The performance for each stimulus pair in the current trial is improved and no modulation by the previous stimulus pairs can be observed. The colorbar corresponds to the fraction classified as *s*_1_ *< s*_2_. **(C)** As in (Fig. 3 B). This improvement of the performance can be traced back to how well the activity bump in the WM network (in purple), before the onset of the second stimulus *s*_2_, tracks the first stimulus *s*_1_ (shown in colored dots, each corresponding to a different value of the inter-stimulus delay interval). Relative to the case in which inputs from the PPC are intact (Fig. 3 A), it can be seen that the location of the bump tracks the first stimulus with high fidelity. The activity in the PPC (in pink), instead, is identical to that shown previously (Fig. 6 A), as all the other parameters are kept constant. **(D)** As in (Fig. 2 C). The bump location can be quantified not only for the stimulus *s*_1_ of the current trial (colored dots, each color corresponding to a given delay interval), but for the four preceding stimuli from the two previous trials (from 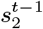 back to 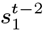). With weaker inputs from the PPC (pink), the WM (purple) function of the circuit is disrupted less frequently, and in the majority of the trials, the bump of activity corresponds to the first stimulus 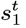.

**Figure S4:**
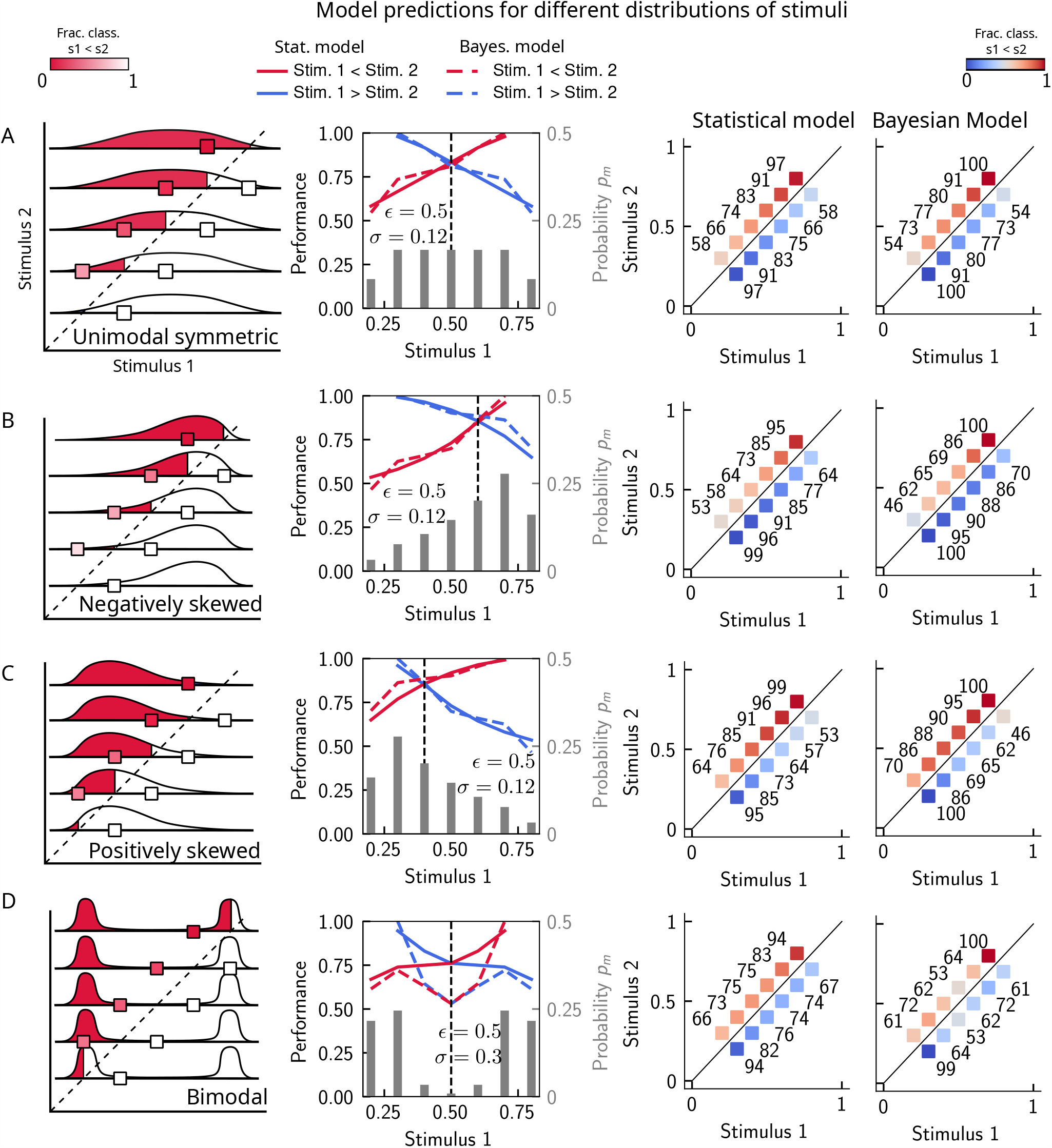
The stimulus distribution impacts the pattern of contraction bias. The model makes different predictions for the performance, depending on the shape of the stimulus distribution. **(A)** Panel 1: schema of model prediction. Regions shaded in red correspond to the probability of correct comparison, for stimulus pairs above the diagonal, when replacing *s*_1_ with a random value sampled from the marginal distribution with a resampling probability *ϵ* = 0.25 (see Fig. 4). Panel 2: prediction of both models for a unimodal symmetric (in this case quasi-uniform) stimulus distribution, statistical model (solid line) and Bayesian model (dashed line). The marginal stimulus distribution is shown in grey bars (to be read with the right y-axis). The value of *s*_1_ for which there is equal performance for pairs of stimuli below and above the diagonal is indicated by the vertical dashed line, corresponding to the median of the distribution. Panel 3: for each stimulus pair, fraction of trials classified as *s*_1_ *< s*_2_ (colorbar), for statistical model. Panel 4: same as panel 3, but for Bayesian model of equal average performance (corresponding to a width of the likelihood of *σ* = 0.08 (see Sect. 1.6.1 and Sect. 4.3). **(B)** Similar to A, for a negatively skewed distribution. **(C)** Similar to A, for a positively skewed distribution. **(D)** Similar to A, for a bimodal distribution.

**Figure S5:**
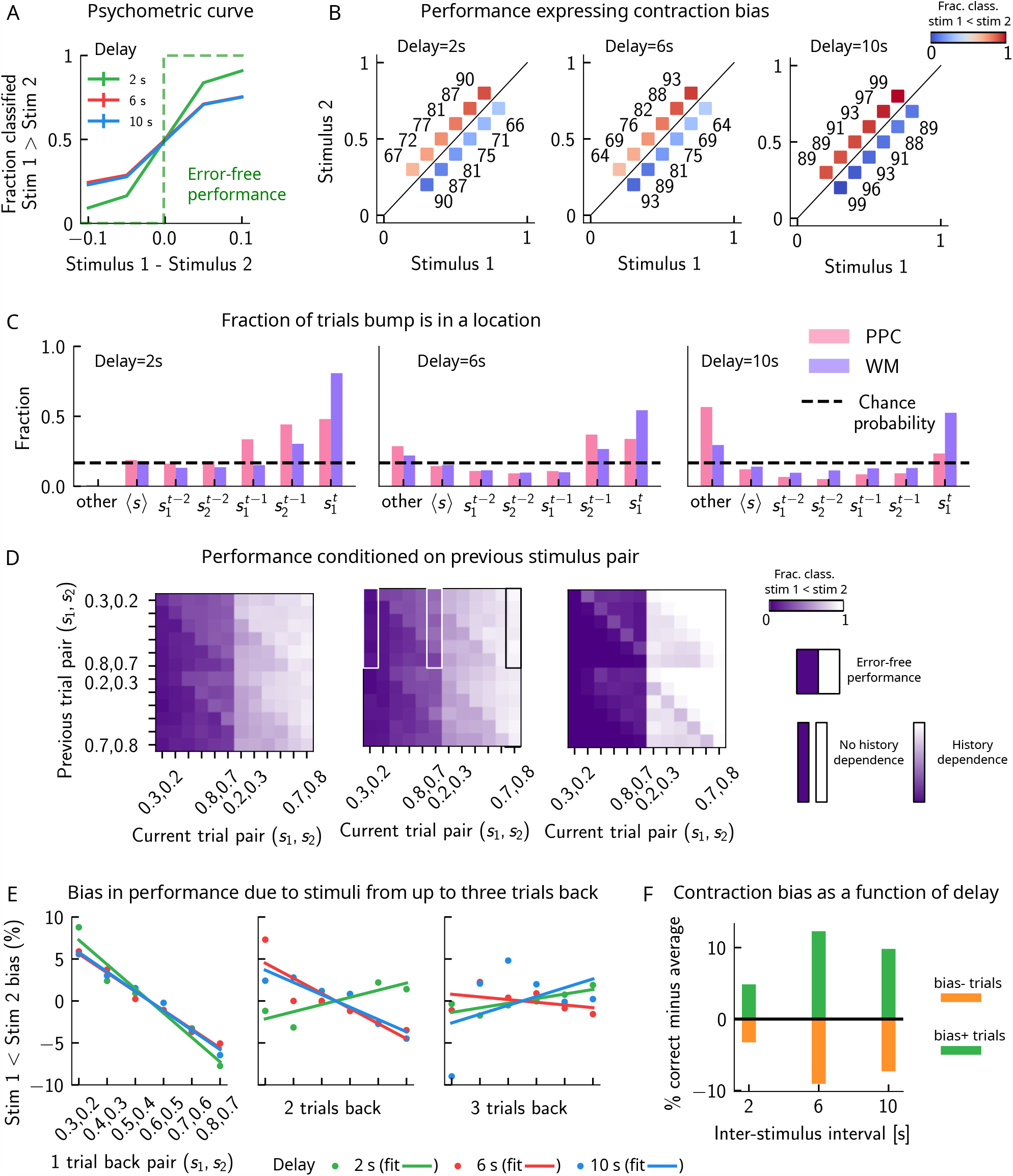
Model predictions for a block design. **(A)** As in (Fig. 2 A). Performance of the network model for the psychometric stimuli improves with a short delay interval and worsens as this delay is increased. **(B)** As in (Fig. 2 C). Performance is affected by contraction bias – a gradual accumulation of errors for stimuli below (above) the diagonal upon increasing (decreasing) *s*_1_. As the delay interval increases, the contraction bias is increased which results in reduced performance across all pairs. Colorbar indicates the fraction of trials classified as *s*_1_ *< s*_2_. **(C)** As in (Fig. 3 C). The location of the bump that corresponds to the value of *s*_1_ occupies a smaller fraction of trials, as the delay interval increases. **(D)** As in (Fig. 2 D). Performance is affected by the previous stimulus pairs (modulation along the y-axis), and becomes worse as the delay interval is increased. The colorbar corresponds to the fraction classified *s*_1_ *< s*_2_. **(E)** As in (Fig. 3 F). Bias, quantifying the (attractive) effect of the previous stimulus pairs, each color corresponding to a different delay interval. These history effects are attractive: the larger the previous trial stimulus pair, the higher the probability of classifying the first stimulus *s*_1_ as large, and vice-versa. Middle/right panels: same as the left panel, for stimuli extending two and three trials back. **(F)** Quantifying contraction bias separately for Bias+ trials (green) and Bias-trials (orange) yields an increasing bias as the inter-stimulus interval increases.

**Figure S6:**
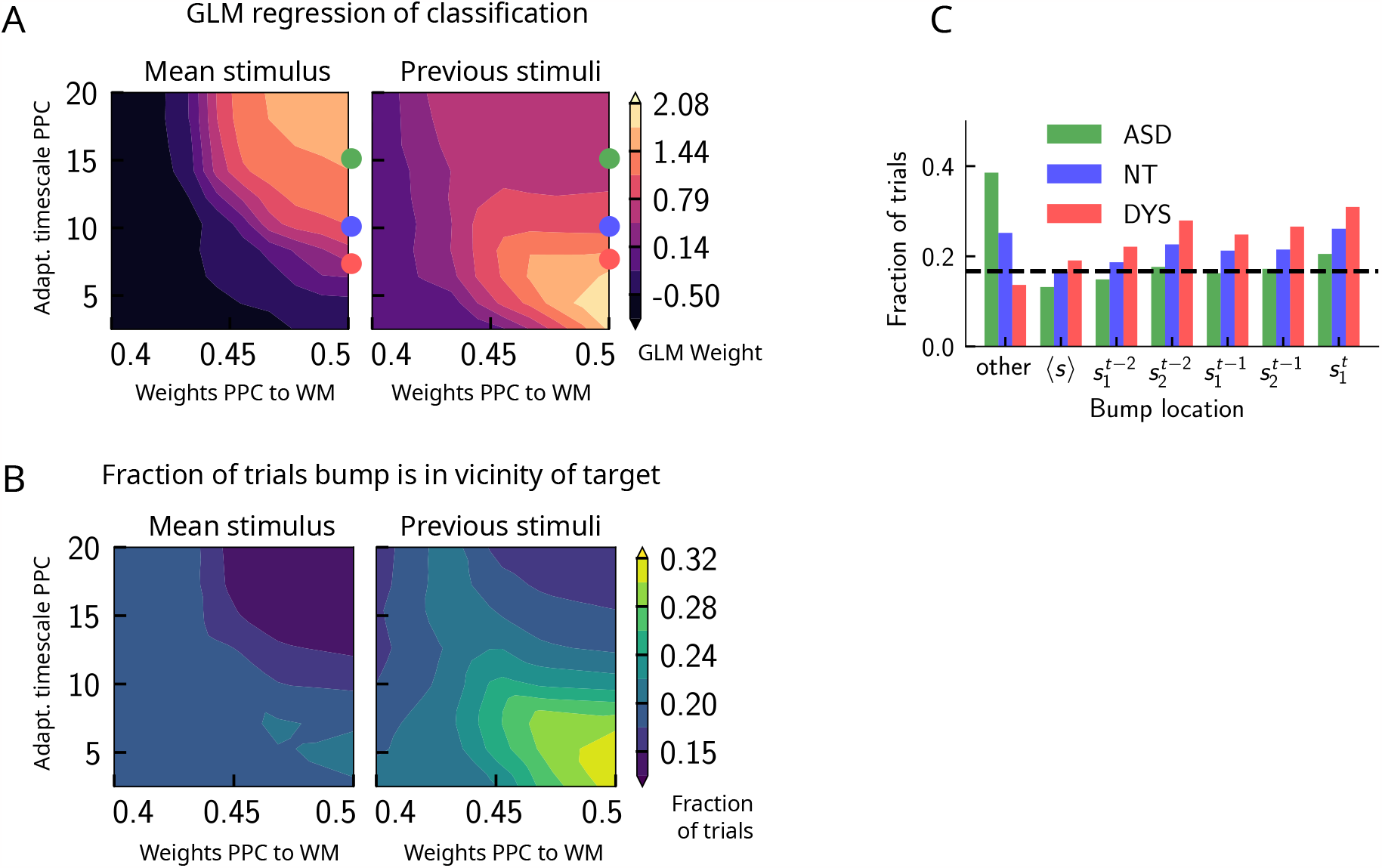
Apparent trade-off between short- and long-term biases, controlled by the timescale of neural adaptation. **(A)** Left: GLM weight associated with the regressor corresponding to the mean stimulus across trials (value indicated by colorbar), as a function of the strength of the weights from the PPC to the WM network (x-axis), and the adaptation timescale in the PPC (y-axis). Right: Same as left panel, but displaying the GLM weight associated with the regressor corresponding to the previous trial’s stimulus. These two panels indicate that the adaptation timescale *seemingly* exerts a trade-off between the two biases: while decreasing it increases short-term sensory history biases, increasing it increases long-term sensory history biases. The values of the adaptation parameter marked by the three colored dots (in red, blue and green) can mimic behaviors similar to dyslexic, neurotypical, and autistic spectrum subjects (see also Fig. 9). **(B)** Left: phase diagram of the fraction of trials in which the activity bump at the end of the delay interval is in the location of the running mean stimulus as a function of the strength of the weights from the PPC to the WM network (x-axis), and the adaptation timescale in the PPC (y-axis). Right: Same as (left), but for the location of any of the two stimuli presented in the previous trial. **(C)** The fraction of trials in which the activity bump at the end of the delay interval corresponds to different locations shown in the x-axis, for three different values of the adaptation timescale parameter, corresponding to qualitatively similar to dyslexic, neurotypical, and autistic spectrum subjects, shown in colors.

### 4.6 Parameters

**Table 2:** Simulation parameters Fig. S1.

| Parameter | Symbol | Default value |
| --- | --- | --- |
| Number of neurons | $N$ | 1000 |
| Neuronal gain | $\beta$ | 5 |
| Range of excitatory interactions [in units of stimulus space length] | $d_0$ | 0.02 |
| Strength of inhibitory weights | $J_0$ | 0.2 |
| Strength of excitatory weights | $J_e$ | 1 |
| Time scale of neuronal integration [s] | $\tau^P$ | 0.01 |
| Amplitude of external inputs | $I_{\text{ext}}$ | 1 |
| Duration of stimuli [s] |  | 0.4 |
| Width of box stimulus [in units of stimulus space length] | $\delta s$ | 0.05 |

**Table 3:** Simulation parameters Fig. 9 and Fig. S6. Other parameters as in Tab. 1.

| Parameter | Symbol | Default value |
| --- | --- | --- |
| Time-scale of neuronal adaptation in PPC [s] | $\tau_\theta^P$ | 7.5 (DYS), 10 (NT), 15 (ASD) |
| Amplitude of adaptation in PPC | $D^P$ | 0.2 |
| Delay interval [s] |  | [2, 6, 10] |
| Inter-trial interval [s] |  | 2.2 |

## Acknowledgements

We are grateful to Loreen Hertäg for helpful comments on our figures, and Arash Fassihi for helpful discussions. We also thank Guilhem Ibos for pointing out a typo in our figure legends in a previous version of this manuscript. This work was supported by BBSRC BB/N013956/1, BB/N019008/1, Wellcome Trust 200790/Z/16/Z, Simons Foundation 564408, EPSRC EP/R035806/1, Gatsby Charitable Foundation 562980 and Wellcome Trust 562763.

